# Serotonin Neurons in the Dorsal and Medial Raphe Nuclei: from Single-Cell Transcriptomes to Whole-Brain Projections

**DOI:** 10.1101/674697

**Authors:** Jing Ren, Alina Isakova, Drew Friedmann, Jiawei Zeng, Sophie Grutzner, Albert Pun, Grace Q. Zhao, Sai Saroja Kolluru, Ruiyu Wang, Rui Lin, Pengcheng Li, Anan Li, Jennifer L. Raymond, Qingming Luo, Minmin Luo, Stephen R. Quake, Liqun Luo

## Abstract

Serotonin neurons of the dorsal and medial raphe nuclei (DR and MR) collectively innervate the entire forebrain and midbrain, modulating diverse physiology and behavior. To gain a fundamental understanding of their molecular heterogeneity, we used plate-based single-cell RNA-sequencing to generate a comprehensive dataset comprising eleven transcriptomically distinct serotonin neuron clusters. Systematic *in situ* hybridization mapped specific clusters to the principal DR, caudal DR, or MR. These transcriptomic clusters differentially express a rich repertoire of neuropeptides, receptors, ion channels, and transcription factors. We generated novel intersectional viral-genetic tools to access specific subpopulations. Whole-brain axonal projection mapping revealed that DR serotonin neurons co-expressing vesicular glutamate transporter-3 preferentially innervate the cortex, whereas those co-expressing thyrotropin-releasing hormone innervate subcortical regions in particular the hypothalamus. Reconstruction of 50 individual DR serotonin neurons revealed segregated axonal projection patterns at the single-cell level. Together, these results provide a molecular foundation of the heterogenous serotonin neuronal phenotypes.

## Introduction

Serotonin is a phylogenetically ancient signaling molecule (Hay-Schmidt, 2000) and the most widely distributed neuromodulator in the brain (Dahlstrom and Fuxe, 1964; Steinbusch, 1981). The serotonin system innervates nearly every region of the brain (Jacobs and Azmitia, 1992), even though it only constitutes ∼ 1/200000 of all CNS neurons in humans. It is critically involved in a broad range of brain functions and is the most frequently targeted neural system pharmacologically for treating psychiatric disorders (Belmaker and Agam, 2008; Ravindran and Stein, 2010).

Serotonin neurons in the central nervous system are spatially clustered in the brainstem, originally designated as groups B1–B9 (Dahlstrom and Fuxe, 1964). Groups B1–B3 are located in the medulla and provide descending serotonergic innervation to the spinal cord and other parts of the medulla. The rest of the groups are located in the pons and midbrain, including the dorsal raphe (DR; groups B6 and B7) and median raphe (MR; groups B5 and B8) nucleus, and provide ascending innervation to the forebrain and midbrain. The DR and MR serotonin systems have been linked with the regulation of many mental states and processes, including anxiety, mood, impulsivity, aggression, learning, reward, social interaction, and hence remain the focus of intense research.

Evidence has suggested that the DR and MR serotonin systems differ in developmental origin, connectivity, physiology, and behavioral function (Calizo et al., 2011; Okaty et al., 2019). DR serotonin neurons derive entirely from rhombomere 1 of the developing mouse brain, whereas MR serotonin neurons derive predominantly from rhombomeres 1, 2, and 3 (Bang et al., 2012; Jensen et al., 2008). Although the DR and MR receive similar inputs globally from specific brain regions (Ogawa et al., 2014; Pollak Dorocic et al., 2014; Weissbourd et al., 2014), they project to largely complementary forebrain targets. The MR serotonin neurons project to structures near the midline, whereas the DR serotonin neurons target more lateral regions (Jacobs and Azmitia, 1992). Slice physiology recording showed that the serotonin neurons in the MR and DR have different electrophysiological characteristics, such as resting potential, resistance, and reaction to serotonin receptor-1A agonist (Calizo et al., 2011). Finally, activation of these two raphe nuclei has been suggested to mediate opposing roles in emotional regulation (Teissier et al., 2015).

Even within the MR or DR, there is considerable heterogeneity of serotonin neurons in multiple aspects. Although MR serotonin neurons arising from different cell lineages are anatomically mixed in the adult, they have distinct electrophysiological properties (Okaty et al., 2015) and potentially distinct behavioral functions (Kim et al., 2009; Okaty et al., 2015). Diversity of serotonin neurons in the DR has received particular attention in recent years. Accumulating evidence indicates that there are subgroup-specific projection patterns within the DR serotonin system (Niederkofler et al., 2016; Ren et al., 2018). The electrophysiological properties of DR serotonin neurons vary according to the projection patterns (Fernandez et al., 2016). Physiological recordings as well as optogenetic and chemogenetic manipulations suggest heterogeneity of DR serotonin neurons in their behavioral functions (Cohen et al., 2015; Marcinkiewcz et al., 2016; Niederkofler et al., 2016). As a specific example, we recently found that DR serotonin neurons that project to frontal cortex and amygdala constitute two sub-systems with distinct cell body locations, axonal collateralization patterns, biased inputs, physiological response properties, and behavioral functions (Ren et al., 2018). Our collateralization analyses also imply that there must be additional parallel sub-systems of DR serotonin neurons that project to brain regions not visited by the frontal cortex- and amygdala-projecting sub-systems.

Ultimately, the heterogeneity of DR and MR serotonin neurons must be reflected at the molecular level. Pioneering work has introduced the molecular diversity of serotonin neurons across the midbrain and hindbrain (Okaty et al., 2015; Spaethling et al., 2014; Wylie et al., 2010), yet systematic analysis and integration of multiple cellular characteristics at the single-cell resolution within each raphe nucleus is still lacking. The rapid development of single-cell RNA sequencing (scRNA-seq) technology in recent years has provided a powerful tool for unbiased identification of transcriptomic cell types in the brain (Darmanis et al., 2015; Li et al., 2017a; Mickelsen et al., 2019; Rosenberg et al., 2018; Saunders et al., 2018; Tasic et al., 2016; Tasic et al., 2018; Welch et al., 2019; Zeisel et al., 2018; Zeisel et al., 2015). In neural circuits where cell types have been well studied by anatomical and physiological methods, there is an excellent correspondence between cell types defined by transcriptomes and the classical methods (Li et al., 2017a; Shekhar et al., 2016). Here, we combine scRNA-seq, fluorescence in situ hybridization, intersectional labeling of genetic defined cell types, whole-brain axonal projection mapping, and single neuron reconstruction to investigate the relationship between molecular architecture of serotonin neurons, the spatial location of their cell bodies in the DR and MR, and their axonal arborization patterns in the brain.

## Results

### Single-cell RNA-sequencing defines 11 transcriptomic clusters of serotonin neurons in the dorsal and medial raphe

We performed a comprehensive survey of DR and MR serotonin neurons in the adult mouse brain by scRNA-seq (***Figure 1A***). To specifically label serotonin neurons, we crossed *Sert-Cre* mice (Gong et al., 2007) with the tdTomato Cre reporter mouse, *Ai14* (Madisen et al., 2010). (Serotonin transporter, or Sert, is a marker for serotonin neurons; see more details below.) We collected serotonin neurons acutely dissociated from brain slices by fluorescence-activated cell sorting (FACS) and used Smart-seq2 (Picelli et al., 2013) to generate scRNA-seq libraries. We used both male and female adult mice (postnatal day 40–45) and applied two dissection strategies to separate the serotonin neurons originating from anatomically-distinct brain regions: 1) in the first set of experiments, we dissected the brainstem region that contain the entire MR and DR; 2) in the second set of experiments, we focused on the principal DR (pDR, corresponding to the traditional B7 group) region by dissecting specifically the DR but excluding its caudal extension (cDR, corresponding to the traditional group B6) (***Figure 1—figure supplement 1A***). After quality control (**Materials and methods**), we determined the transcriptomes of 709 cells from eight samples that include MR, pDR, and cDR, and 290 cells from six pDR-only samples (999 cells in total). We sequenced to a depth of ~1 million reads per cell and detected ∼10,000 genes per cell (***Figure 1—figure supplement 1B***, ***Supplemental Table 1***). The data quality and serotonin identity were validated by the fact that all 999 neurons expressed: 1) tryptophan hydroxylase 2 (Tph2), a key enzyme for serotonin biosynthesis; 2) transcription factor Pet1 (Fev) known to express in raphe serotonin neurons (Hendricks et al., 1999); 3) the plasma membrane serotonin transporter (Sert), which recycles released serotonin back to presynaptic terminals of serotonin neurons, and 4) vesicular monoamine transporter 2 (Vmat2), which transports serotonin (and other monoamines) from presynaptic cytoplasm to synaptic vesicles (***Figure 1—figure supplement 1C***).

**Figure 1.**
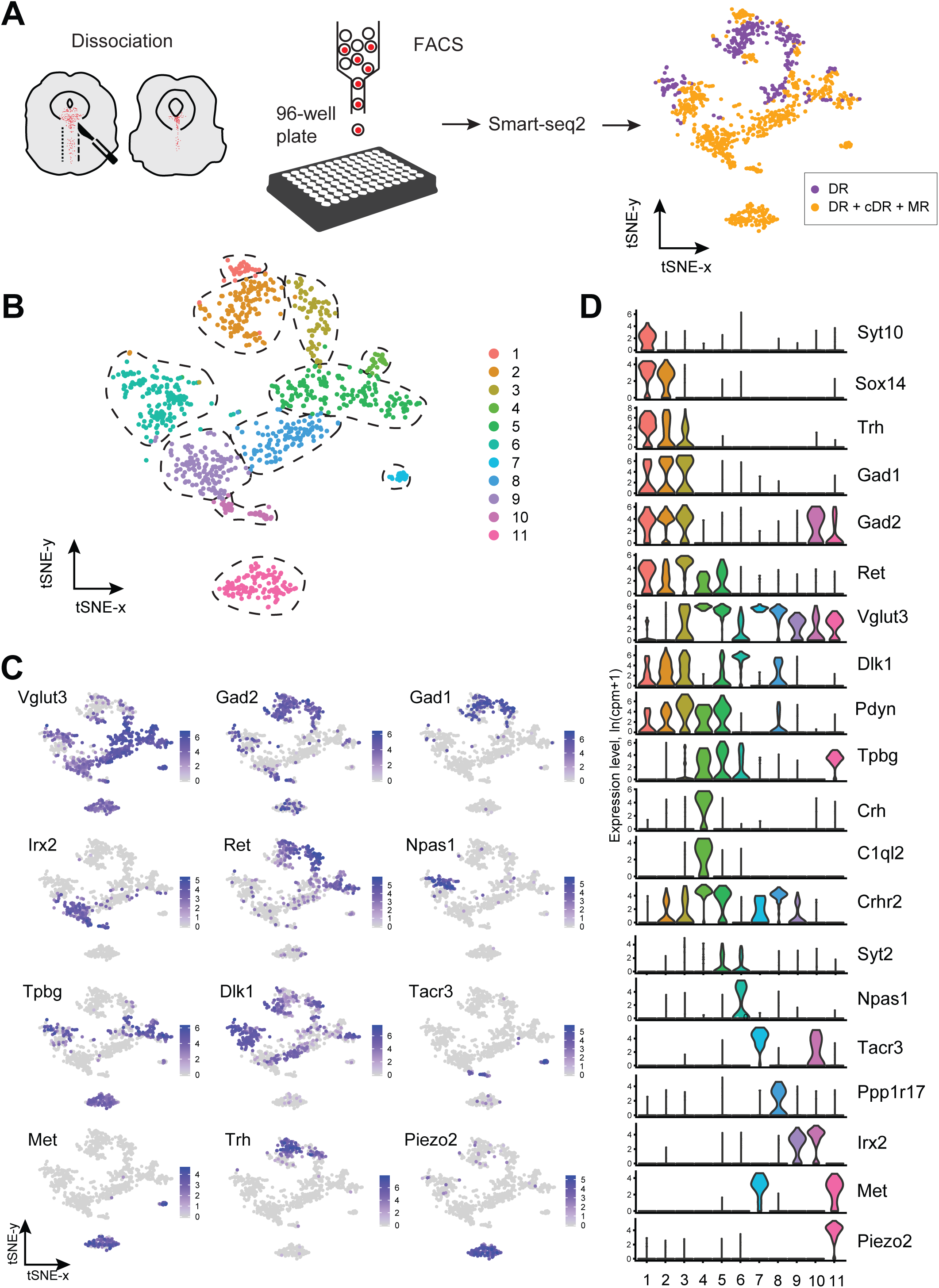
Single-cell transcriptomic profiling of serotonergic neurons. **(A)** Schematic representation of scRNA-seq pipeline used to analyze individual serotonin neurons. Tissue slices containing either principal dorsal raphe (pDR, n = 6 brains) or all raphe nuclei (n = 8 brains) of C57BL/6J adult were dissociated to a single-cell suspension. tdTomato*^+^* neurons were FACS-sorted in 96-well plates and processed for scRNA-seq using Smart-seq2 protocol. tSNE plot of all processed *Tph2^+^* neurons colored by anatomical localization. cDR, caudal DR; MR, medial raphe. **(B)** tSNE plot of 999 *Tph2^+^* cells obtained from 14 brains and clustered by gene expression. Cells are colored-coded according to identified transcriptomic clusters. **(C)** Expression of genes defining distinct serotonin neuron populations. Cells are colored by log-normalized expression of each transcript, and the color legend reflects the expression values of ln(CPM+1). CPM, counts per million. **(D)** Violin plots of expression of marker genes across 11 clusters.

To define distinct serotonin neuron populations based on single-cell transcriptome, we performed principal component analysis (PCA) on all the genes expressed in the assayed neurons, followed by nearest-neighbor graph-based clustering. These 999 MR and DR serotonin neurons comprised 11 clusters (***Figure 1B***, ***Figure 1—figure supplement 2*** and **Materials and methods**). Each cluster contained cells from both sexes after removing genes located on the Y chromosome, indicating that there were few sex-specific differences (***Figure 1—figure supplement 1D***). No substantial batch effect was observed (***Figure 1—figure supplement 1E***). Each of the 11 clusters expressed a set of cluster-discriminatory genes, including markers for specific neurotransmitter systems, such as *Vglut3*, *Gad1*, and *Gad2* (***Figure 1C,D; Figure 1—figures supplement 3–6***).

### Anatomical organization of transcriptomically defined serotonin clusters

Of the 11 transcriptomic clusters, six (Cluster 1–6) consisted of only serotonin neurons dissected from the pDR. We hypothesized that these six clusters represent serotonin neurons from the pDR, and the remaining five clusters represent cells from MR and cDR. To test this hypothesis and to obtain information about the anatomical organization of transcriptomically defined serotonin cell clusters within the DR and MR, we chose 16 cluster marker genes and performed hybridization-chain reaction (HCR)-based single-molecule fluorescence in situ hybridization (smFISH) (Choi et al., 2018). To restrict our analysis within the serotonin neuron population, we simultaneously double-labeled *Tph2*, a marker for serotonin neurons, for all the HCR-smFISH experiments.

***Figure 2A*** summarizes the distribution of the *Tph2^+^* serotonin neurons that express each of the 16 cluster markers in four coronal sections that cover the pDR, cDR, and MR. This summary was derived from counting Tph2/cluster marker double-labeled cells from confocal sections of the HCR-smFISH experiments ***(Figure 2B*** and ***Figure 2—supplement figures 1–2***). Specifically, we found that the distribution of markers for Clusters 1–6, *Ret*, *Trh*, *Gad1*, *Npas1*, *Syt2*, and *C1ql2* (***Figure 1D***), were mostly restricted to the pDR. The two common Cluster 7 markers *Tacr3* and *Met* were both highly concentrated in serotonin neurons under the aqueduct in the cDR. *Dlk1* should be expressed in the DR clusters as well as Cluster 8 (***Figure 1D***), and its expression was found in both the DR and MR, suggesting Custer 8 serotonin neurons are located in the MR. Clusters 9–11 markers *Irx2* and *Piezo2* were mostly found in the MR. Thus, these observations support the anatomical breakdown suggested by the dissection of primary tissue, and additionally provide a more granular and detailed description about finer boundaries.

**Figure 2.**
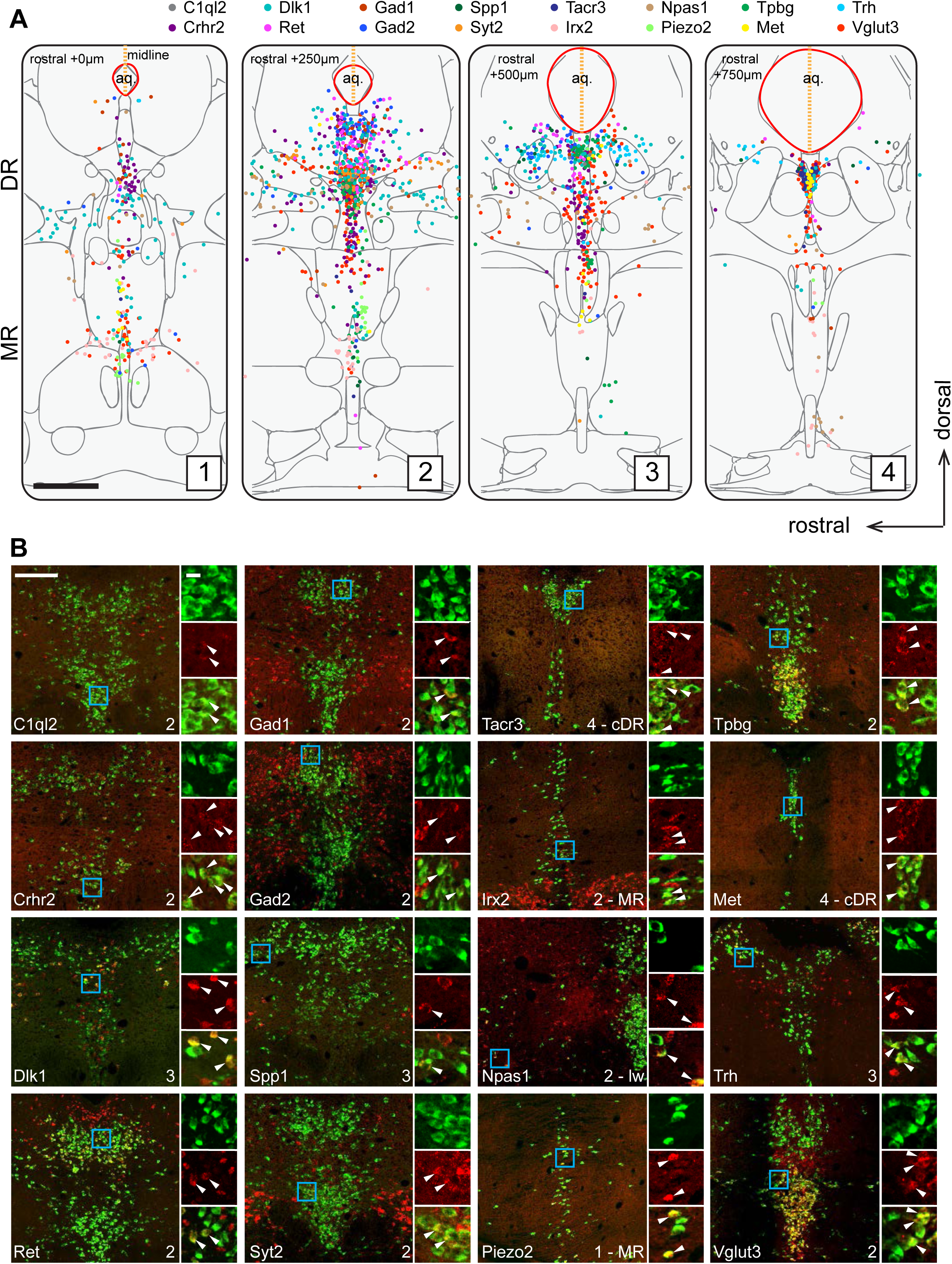
Anatomical location of serotonin neuron clusters determined by hybridization-chain reaction-based single-molecule fluorescence in situ hybridization (HCR-smFISH) of 16 cluster markers. **(A)** Positions of double-positive neurons (*Tph2* and each of 16 marker genes color-coded on the top) are shown on 4 schematics representing coronal slices 250 µm apart. Red line around the aqueduct represents the average boundary drawn from raw data for each slice. Scale bar, 500 µm. **(B)** Representative images for each of the genes schematized in (A). All green cells are Tph2-positive, red cells express the indicated marker gene. Each image corresponds to one of the four numbered schematics in (A) and is located immediately ventral to the aqueduct unless otherwise noted as MR, lateral wing (lw), or cDR. Cyan box highlights the individual color zoom region at right. White arrowheads mark examples of double-positive neurons. Scale bars, 200 µm main panels, 20 µm zoom panels.

Within the DR, *Trh^+^*, *Gad1^+^*, and *Gad2^+^* serotonin neurons were mainly located in the dorsal DR, whereas *Vglut3^+^* and *Syt2^+^* serotonin neurons were mainly located in the ventral DR and cDR. These data suggest that Clusters 1–3 correspond to the dorsal DR and Cluster 4–6 to the ventral DR. Cluster 6 marker *Npas1^+^* was largely excluded from the densest portion of *Tph2* expression at the midline and instead was found scattered in the more rostral and ventral portion of the lateral wings. On the other hand, *Crhr2*, which should be expressed in all DR serotonin neuron clusters except Cluster 1 and 6 (***Figure 1D***), was localized preferentially near the midline and was absent from the lateral wing. Thus, Cluster 6 likely corresponds to serotonin neurons located preferentially in the lateral wings. In contrast to DR, the anatomical organization of the molecular features that define MR clusters is less obvious and different clusters appear more intermingled.

In summary, our HCR-smFISH experiments support the notion that Clusters 1–6 correspond to pDR serotonin neurons, Cluster 7 corresponds to cDR serotonin neurons, and Clusters 8–11 correspond to MR serotonin neurons. We thus rename hereafter Clusters 1–6 as DR-1–6, Cluster 7 as cDR, and Clusters 8–11 as MR-1–4 (***Figure 3***). Detailed expression levels of marker genes across all 11 clusters can be found in ***Figure 3—figure supplement 1–3***.

**Figure 3.**
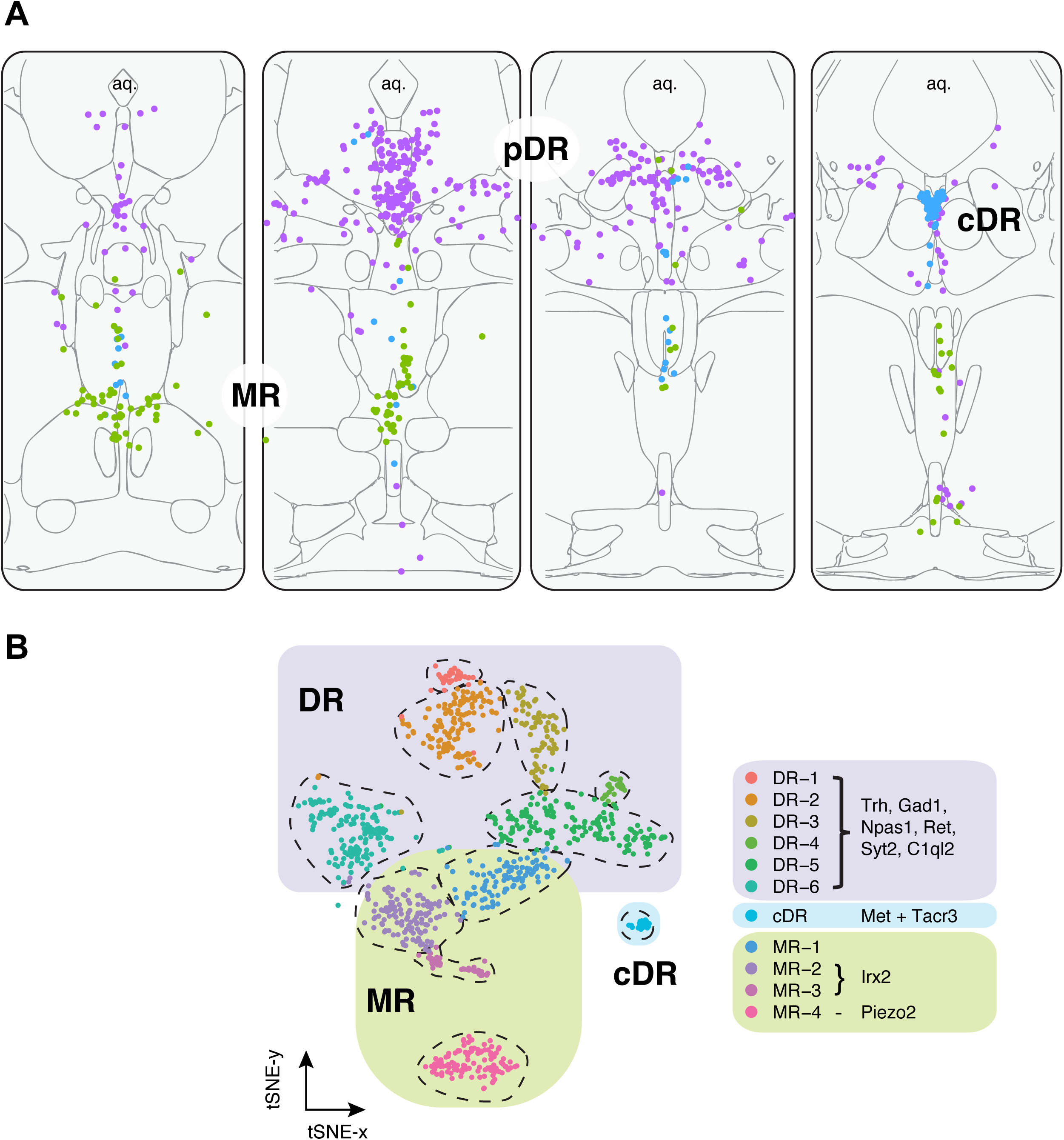
Summary of spatial distribution of transcriptomic clusters of serotonin neurons. **(A)** Purple dots represent distribution of Clusters 1–6 markers listed in panel (B); green dots represent the distribution *Irx2* and *Piezo2* cells that are Cluster 9–11 markers; and cyan dots present distribution of *Met^+^* and *Tacr3^+^* cells, which are co-expressed in Cluster 7, but is additionally expressed in Cluster 10 (*Met*) or Cluster 11 (*Tacr3*). **(B)** Collectively, scRNA-seq and HCR-smFISH experiments support the model that Clusters 1–6 from Figure 1B correspond to pDR serotonin neurons (renamed DR-1–6), Cluster 7 corresponds to cDR serotonin neurons, and Clusters 8–11 correspond to MR serotonin neurons.

### Molecular properties of MR and DR serotonin neurons

Having determined the spatial locations of transcriptomically defined serotonin cell types, we next analyzed key groups of differentially expressed genes crucial for neuronal function, including markers for neurotransmitter systems, neuropeptides, ionotropic and metabotropic (G-protein-coupled) neurotransmitter receptors, wiring specificity molecules, and transcription factors (***Figure 4***; ***Figure 4—figure supplement 1***).

**Figure 4.**
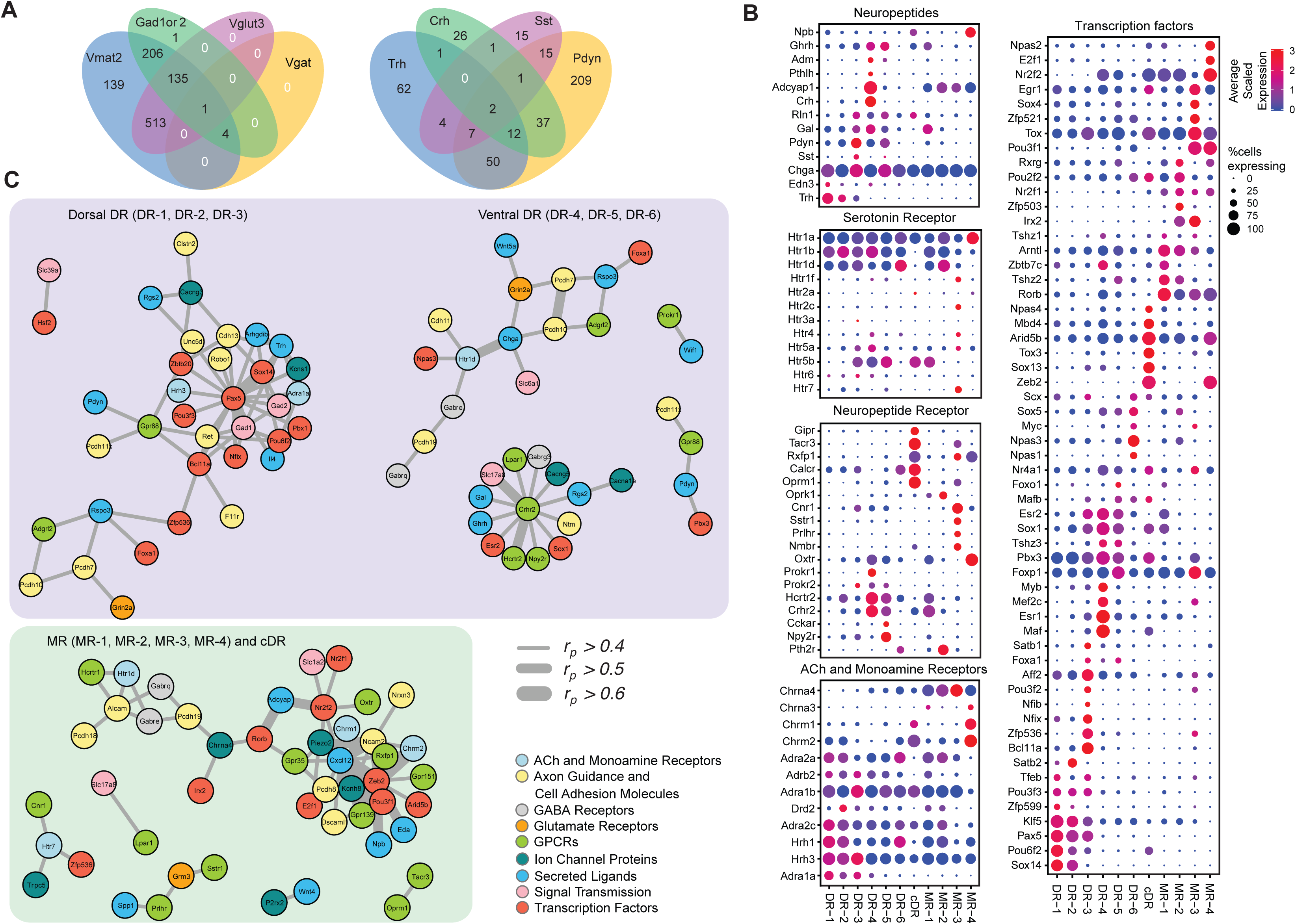
Molecular and functional characteristics of distinct serotonin neuron clusters. **(A)** Venn diagram showing the number of cells co-expressing genes associated with markers for different neurotransmitter systems: *Gad1/2*, *Vmat2*, *Vgat*, and *Vglut3* (left) and neuropeptides: *Trh*, *Crh*, *Sst*, *Pdyn* (right). We consider a gene to be expressed if it has at least one read mapping to it and is detected in at least 3 cells (***Materials and methods***). **(B)** Expression of the most variable neuropeptides, receptors, and transcription factors across molecularly distinct serotonin neuron clusters. **(C)** Network representation of co-expressed genes that belong to one of the functional gene categories that organize transcriptional regulation, synaptic connectivity, and neuronal communications. Networks were constructed based on Pearson correlation coefficient (*r*p) of gene expression across all cells and centered on pDR- and MR+cDR-specific genes. Genes appear connected if *r*p >0.4. Line width represents *r*p. Nodes are colored according to functional gene categories.

#### Genes related to neurotransmitters other than serotonin

The majority of the clusters express *Slc17a8*, which encodes vesicular glutamate transporter-3 (Vglut3). These include almost all serotonin neurons from the cDR and the majority from MR clusters, and DR-3–6 clusters (***Figure 1D***). These observations suggest that glutamate is the most prevalent co-transmitter for serotonin neurons. Glutamate co-release from Vglut3^+^ serotonin terminals has indeed been reported at the orbital prefrontal cortex (Ren et al., 2018), nucleus accumbens (Liu et al., 2014), ventral tegmental area (Wang et al., 2019), and basolateral amygdala (Sengupta et al., 2017).

Three DR clusters express *Gad1* and *Gad2*, which encode biosynthetic enzymes for the neurotransmitter GABA. Two MR clusters express *Gad2* but not *Gad1*, and cDR expresses neither. Interestingly, few of the *Gad1^+^* or *Gad2^+^* neurons express vesicular GABA transporter (*Vgat*, ***Figure 4A*** left), which is responsible for transporting GABA into synaptic vesicles for synaptic transmission. However, it has been reported that vesicular monoamine transporters (*Slc8a1* for Vmat1; *Slc8a2* for Vmat2) can transport GABA into synaptic vesicles (Stensrud et al., 2014), and virtually all serotonin neurons expressed Vmat2 (***Figure 1—figure supplement 1D***, ***Figure 4A*** left). Nevertheless, it remains to be determined if these serotonin neurons can actually release GABA. At the single cell level, 15% of serotonin neurons do not express any of the gene markers for glutamate or GABA transmission. 50% of serotonin neurons express *Vglut3*, 21% express either *Gad1*, *Gad2*, or both, and 13% express markers for *Vglut3* and *Gad1* or *Gad2* (***Figure 4A*** left).

In addition to small-molecule transmitters, the majority of serotonin neurons also co-express neuropeptides (***Figure 4A*** right). Expression of several neuropeptides served as excellent cluster markers. For example, thyrotropin-releasing hormone (Trh) is highly expressed in DR-1–3 (14% serotonin neurons, ***Figure 1C***). Corticotropin-releasing hormone (Crh) is highly expressed in the DR3 cluster but much less everywhere else (7% serotonin neurons). Neuropeptide B (Npb) is highly expressed in cDR and MR4 but much less in pDR serotonin neurons. Many serotonin neurons express multiple neuropeptides (***Figure 4A,B***).

#### Small-molecule neurotransmitter receptors

Each cluster has a distinct expression pattern of neurotransmitter (including neuropeptide) receptors (***Figure 4B***, ***Figure 4—figure supplement 1***). Multiple subunits of glutamate and GABA receptors are differentially expressed across the clusters. For example, *Grin3a*, encoding subunit 3A of ionotropic NMDA glutamate receptor, is mainly expressed in pDR but not MR or cDR clusters. By contrast, *Grik1*, encoding subunit 1 of the kainate glutamate receptor, is more highly expressed in MR clusters. *Chrna3* and *Chrna4*, encoding subunit α3 and α4 of the nicotinic acetylcholine receptor, are also more enriched in MR clusters. Clusters MR-4 and cDR have the highest expression level of expression for *Chrm1* and *Chrm2*, encoding muscarinic acetylcholine receptors. Finally, all serotonin neurons express at least one type of serotonin receptors, with the Gi-coupled *Htr1* subfamily in particular *Htr1a* being the most widely expressed. *Htr1a* is highly expressed in the MR-4 cluster, which does not express other serotonin receptors. Of all the genes encoding serotonin receptors, *Htr5b* has the strongest cluster specificity (***Figure 4B***).

##### Neuropeptide receptors

Multiple neuropeptide receptors have rich expression in distinct clusters. Notably, cDR preferentially expresses at least five neuropeptide receptors: gastric inhibitory polypeptide receptor (*Gipr*), tachykinin receptor 3 (*Tacr3*), relaxin family peptide receptor 1 (*Rxfp1*), calcitonin receptor (*Calcr*), and opioid receptor mu-1 (*Oprm1*). Meanwhile, opioid receptor kappa 1 (*Oprk1*) is expressed specifically in MR-2. Another neuropeptide receptor enriched cluster is MR-3, expressing *Tacr3*, cannabinoid receptor 1(*Cnr1*), somatostatin receptor-1 (*Sstr1*), prolactin-releasing hormone (*Prlhr*), and neuromedin B receptor (*Nmbr*). Neuropeptide receptors like oxytocin receptor (*Oxtr*), prokineticin receptor-1 and −2 (*Prokr1*, *Prokr2*), hypocretin receptor-2 (*Hcrtr2*), corticotropin-releasing hormone receptor-2 (*Crhr2*), cholecystokinin-A (*Cckar*), neuropeptide Y receptor Y2 (*Npy2r*), and parathyroid hormone-2 receptor (*Pth2r*) all have distinct expression pattern across different clusters.

In summary, the DR and MR serotonin neurons have diverse expression pattern of neuropeptide receptors, indicating that they are subject to complex modulation by a host of neuropeptides.

##### Other notable genes

the DR and MR serotonin clusters are also distinguished by differential expression of ion channels as well as axon guidance and cell adhesion molecules (***Figure 4—figure supplement 1***), which can contribute to their differences in physiological properties and wiring specificity. Notably, genes encoding a voltage-gated K^+^ channel (*Kchn8*) and mechanosensitive channel (*Piezo2*) are highly expressed in the MR-4 cluster but exhibit little expression in all other clusters. The chemokine ligand-12 (*Cxcl12*) and cadherin-6 (*Cdh6*) are preferentially expressed in MR-1 and DR-4 clusters, respectively.

##### Transcription factors

Transcription factors (TFs) have been shown to be the best molecular feature for cell type definition (Tabula Muris et al., 2018). Within our data, we observed robust cluster-specific expression of multiple TF genes (***Figure 4B***; ***Figure 4—figure supplement 1***). Importantly, among the genes we identified are those previously reported to be involved in neuronal function. For example, *Nfix* and *Nfib* (Bedford et al., 1998) are preferentially expressed in DR-3. *Irx2* (Wylie et al., 2010) is specific to MR-2 and MR-3. *Sox13* is highly enriched in cDR. *Zeb2* (Okaty et al., 2015) is uniquely expressed in cDR and MR-4. TFs associated with neurodevelopmental disorders, such as *Npas1* and *Npas3* (Erbel-Sieler et al., 2004), are preferentially expressed in DR-6, and *Aff2 (Mondal et al., 2012)* is enriched within DR-3.

##### Transcriptional networks

To understand the relationship between cluster-specific genes and to gain new insights on transcriptional regulatory programs orchestrating the maintenance of serotonin neuron subtype identity, we next performed a pairwise correlation analysis of gene expression across all 999 neurons (***Supplemental File 1***). We used Pearson correlation coefficient (*rp*) as a measure of gene co-expression and focused on the genes with average expression > 10 counts per million. We found multiple genes to co-expressed (defined as *rp> 0.5*) in serotonin neurons, forming co-expression modules among each other as well as around various TFs (***Supplemental File 1***). Focusing further on cluster markers, we identified three transcriptional “hubs” that putatively govern molecular programs within respective neurons. Interestingly, two of these hubs were centered around pDR-specific TFs, and, independently, comprised of dorsal (DR-1, DR-2, DR-3) and ventral (DR-4, DR-5, DR-6) pDR markers (***Figure 4C***). Among dorsal pDR markers we identified *Pax5* and *Sox14* to strongly correlate (*rp*>0.5) with *Gad1*, *Gad2*, and *Trh*, among other genes. Among ventral pDR markers, we found *Crhr2* to be highly correlated with several TFs, receptors, and neurotransmitter-related genes, most notably *Vglut3* (*Slc17a8*) and *Hcrtr2*. These *Crhr2^+^* serotonin neurons could use TFs *Esr2* and *Sox1* to maintain their subtype identity. *Npas3* is specifically expressed in the DR-6 cluster and is highly correlated with *Htr1d*, suggesting a critical role of maintaining the characteristics of DR-6 serotonin neurons. Similarly, we found several MR and cDR-specific TFs to be the hubs of co-expression modules. Particularly, the expression of a large number of cell adhesion molecules, receptors, ion channel proteins and neuropeptides strongly correlate with the expression of TFs *Zeb2*, *Pou3f1*, *Irx2*, and *Zfp536*. Based on the identified correlation among gene expression of multiple marker genes across all the cells, we speculate that the identified genes are linked by one or multiple transcriptional regulatory programs, which, in turn, drive cell type-specific functions of distinct serotonin neuron populations.

##### Sexually dimorphic gene expression

Finally, even though there were no apparent sex-specific difference at the cluster level ***Figure 1—figure supplement 1D***), we did detect several genes, such as *Sod1*, *Snx10, Inpp4a, Zscan26, Ncam1*, showing sexual dimorphism across the majority of cell subtypes (***Figure 4—figure supplement 2***).

### Viral-genetic tools to access different serotonin neuron subtypes

The gene expression patterns of specific serotonin neuron clusters can in principle allow genetic access to these specific subpopulations for anatomical tracing, physiological recording, and functional perturbation (Luo et al., 2018). However, DR and MR contain not only serotonin neurons but also GABAergic and glutamatergic neurons that do not express Tph2 (and hence do not release serotonin), some of which may project to the same target regions (McDevitt et al., 2014). To precisely investigate the function of transcriptome-based serotonin neuronal types, we need to use an intersectional strategy to target serotonin neurons that express specific additional markers (Jensen et al., 2008). To this end, we generated *Sert-Flp* mice through homologous recombination-based knock-in in embryonic stem cells (**Materials and methods**), and used *Sert-Flp* mice to intersect with transgenic Cre mice that allow expression of a fluorescent reporter only in Flp^+^/Cre^+^ (AND gate), so as to genetically label only specific Cre-positive clusters (***Figure 5A***).

**Figure 5.**
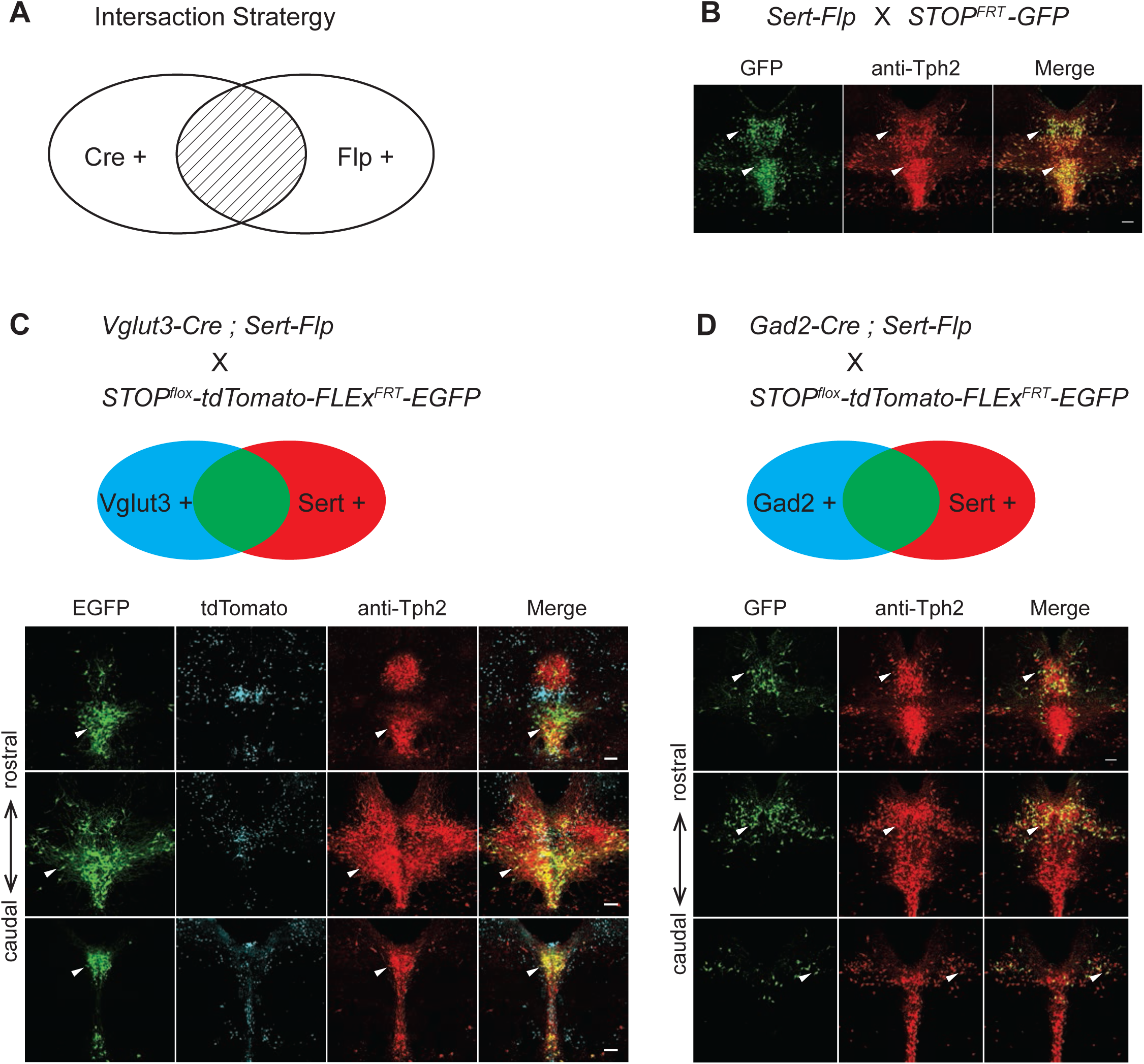
Intersectional strategy to genetically access specific serotonin neuron populations. **(A)** Schematic representing cells dually gated by Cre- and Flp-expression. **(B)** *Sert-Flp* mice were crossed with *H11-CAG-FRT-STOP-FRT-EGFP* (*STOP^FRT^-GFP*) mice. Anti-Tph2 staining (red) was performed on consecutive coronal sections containing DR. 98.5% GFP^+^ neurons are Tph2^+^ and 100% Tph2^+^ neurons are GFP^+^ (n = 3 mice). **(C)** In mice triple transgenic for *Vglut3-Cre*, *Sert-Flp*, and the *IS* reporter (*Rosa-CAG-loxP-stop-loxP-FRT-tdTomato-FRT-EGFP*), EGFP^+^ (Flp^+^Cre^+^) cells are referentially found in ventral pDR and in cDR (arrowheads). Coronal sections containing DR are shown, counterstained with Anti-Tph2 (red). **(D)** In mice triple transgenic for *Gad2-Cre*, *Sert-Flp*, and the *IS* reporter, EGFP^+^ (Flp^+^Cre^+^) cells are referentially found in dorsal pDR (arrows). Coronal sections containing DR are shown, counterstained with Anti-Tph2 (red). Scale, 100 µm.

To characterize the *Sert-Flp* mouse line, we crossed it with *H11-CAG-FRT-stop-FRT-EGFP* mice we newly generated (**Materials and methods**). Anti-Tph2 staining on the brain slices containing pDR showed that 98.5% GFP^+^ neurons are Tph2^+^ and 100% Tph2^+^ neurons are GFP^+^ (***Figure 5B***). To further verify the intersectional strategy and to label the serotonin neurons co-expressing markers for glutamate or GABA transmission, we crossed *Sert-Flp* with the *IS* reporter mice (*Rosa-CAG-loxP-stop-loxP-FRT-tdTomato-FRT-EGFP*) (He et al., 2016) and either *Vglut3-Cre* or *Gad2-Cre*. Anti-Tph2 staining showed that all GFP-labeled neurons are Tph2^+^ (***Figure 5C,D***). In the pDR, Vglut3^+^ serotonin neurons were mainly located ventrally, whereas Gad2^+^ serotonin neurons were mainly located dorsally, consistent with our previous study (Ren et al., 2018) and the HCR-smFISH results (***Figure 2***).

To map the axonal projection pattern of serotonin subtypes defined by intersection of Flp and Cre expression, we developed a new AAV vector (*AAV-CreON/FlpON-mGFP*) that expressed membrane-targeted GFP under the dual gates of Flp and Cre (***Figure 6A***). Based on our scRNA-seq and HCR-smFISH results, the *Vglut3^+^* and *Trh^+^* pDR serotonin neurons consist of largely complementary cell types at the transcriptomic level and have a distinct distribution along the dorsal– ventral axis in the pDR. To visualize these two subpopulations of serotonin neurons, we injected *AAV-CreON/FlpON-mGFP* into pDR of either *Vglut3-Cre;Sert-flp* (n = 3, ***Figure 6B***) or *Trh-Cre;Sert-flp* mice (n = 3, ***Figure 6C***). Anti-Tph2 staining showed that 98.2% GFP^+^ neurons were Tph2^+^. As predicted, *Vglut3^+^Sert^+^* GFP cells were mostly located in the ventral pDR (***Figure 6B***), whereas *Trh^+^Sert^+^* cells were located in the dorsal pDR (***Figure 6C***). As negative controls, we injected the same virus into mice carrying only the *Sert-flp* transgene or only the *Vglut3-Cre* transgene and did not find any mGFP^+^ cell bodies or fibers (n = 3 for each; data not shown).

**Figure 6.**
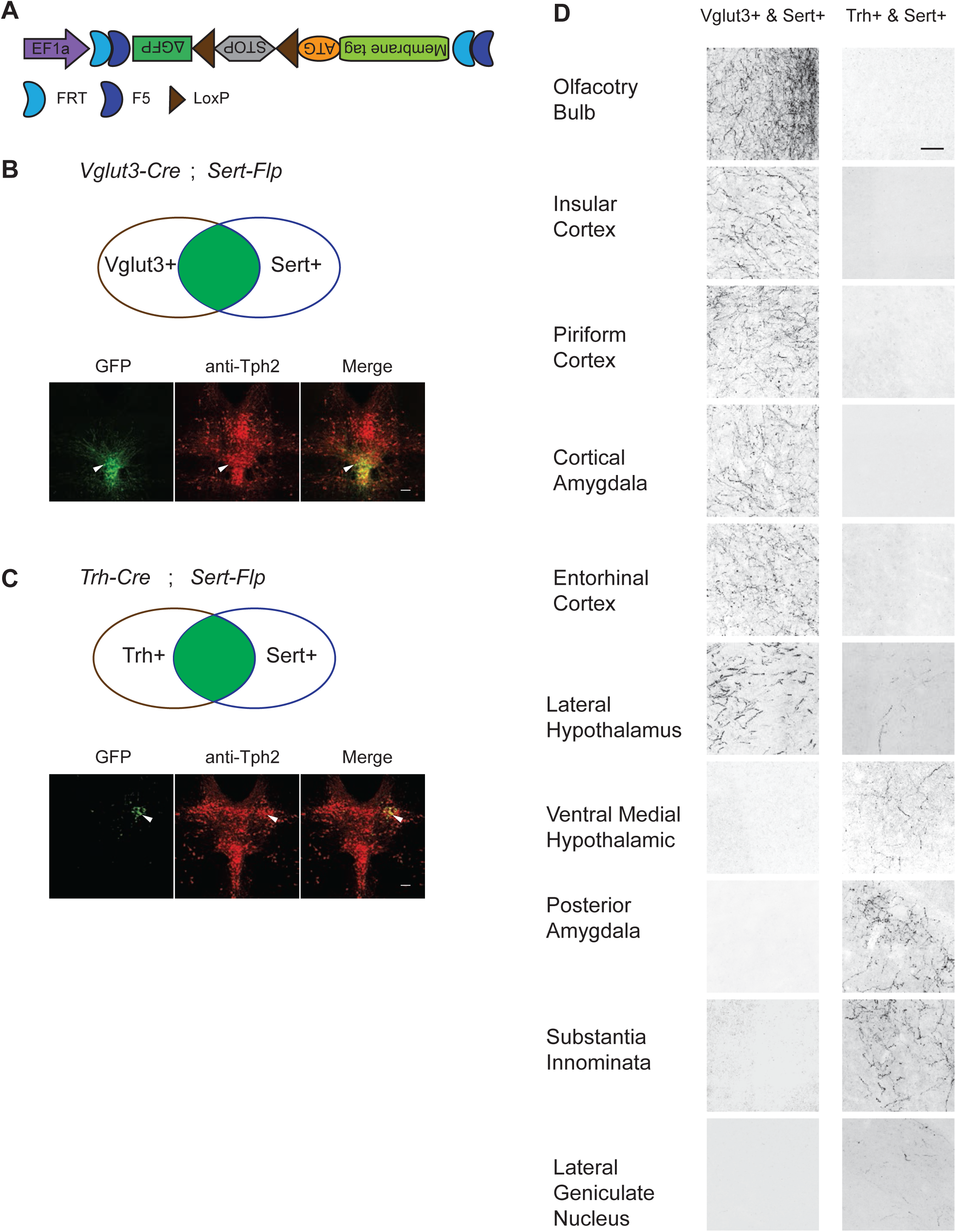
Dually gated serotonin neuron and axonal terminal labeling by viral-genetic intersection. **(A)** Schematics representing of the *AAV-CreON/FlpON-mGFP* dual labeling design. **(B)** mGFP expression (green) and Tph2 immunoreactivity (red) after injection of *AAV-CreON/FlpON-mGFP* into the DR of *Sert-Flp*;*Vglut3-Cre* mice. mGFP is mostly restricted to ventral pDR. **(C)** mGFP expression (green) and Tph2 immunoreactivity (red) after injection of *AAV-CreON/FlpON-mGFP* into the DR of *Sert-Flp*;*Gad2-Cre* mice. mGFP is mostly restricted to dorsal pDR; the left-right asymmetry was likely due to AAV injection being biased towards the right hemisphere. **(D)** Axonal terminal expression of mGFP in different brain regions of *Sert-Flp*;*Vglut3-Cre* mice and *Sert-Flp*;*Gad2-Cre* mice. The left and right images represent comparable brain regions cropped from serial coronal sections. Scale, 100 µm.

The intersectional strategy allowed us to trace the projection of GFP^+^ axons from these two groups of serotonin neurons across the brain. We next examined their projections by staining every four coronal sections across the brain with anti-GFP antibody. We found that serotonin axons from *Vglut3^+^* population preferentially targeted cortical regions (***Figure 6D***), consistent with our previous results (Ren et al., 2018). By contrast, no GFP-labeled axons were observed in the cortical regions from *Trh-Cre:Sert-flp* mice. Instead, *Trh^+^* serotonin axons project to the anterior and medial hypothalamus, posterior amygdala, and the lateral geniculate nucleus in the thalamus, none of which were targeted by *Vglut3^+^* axons (***Figure 6D***).

### Whole-brain axonal projections of selected serotonin neuron subpopulations

While suggestive of an anatomical division in targets, assessing the full extent to which projections of *Vglut3^+^* or *Trh^+^* pDR serotonin neuron populations segregate requires quantifying axonal innervation at the whole-brain level. We used the iDISCO-based brain clearing technique AdipoClear (Chi et al., 2018) to visualize, align, and summarize whole-brain projectomes (***Figure 7A***). Individual hemispheres of either *Vglut3-Cre;Sert-flp* (n = 3) or *Trh-Cre;Sert-flp* mice (n = 3) injected with *AAV-CreON/FlpON-mGFP* at pDR were imaged by light-sheet microscopy. We developed deep learning models to automatically trace whole-brain axonal projections by segmenting volumes with a 3D UNet-based convolutional neural network we developed (**Materials and methods**). The resulting volumetric probability maps were thinned and thresholded before aligning to the Allen Institute’s 2017 common coordinate framework as previously described (Ren et al., 2018; ***Figure 7A***).

**Figure 7.**
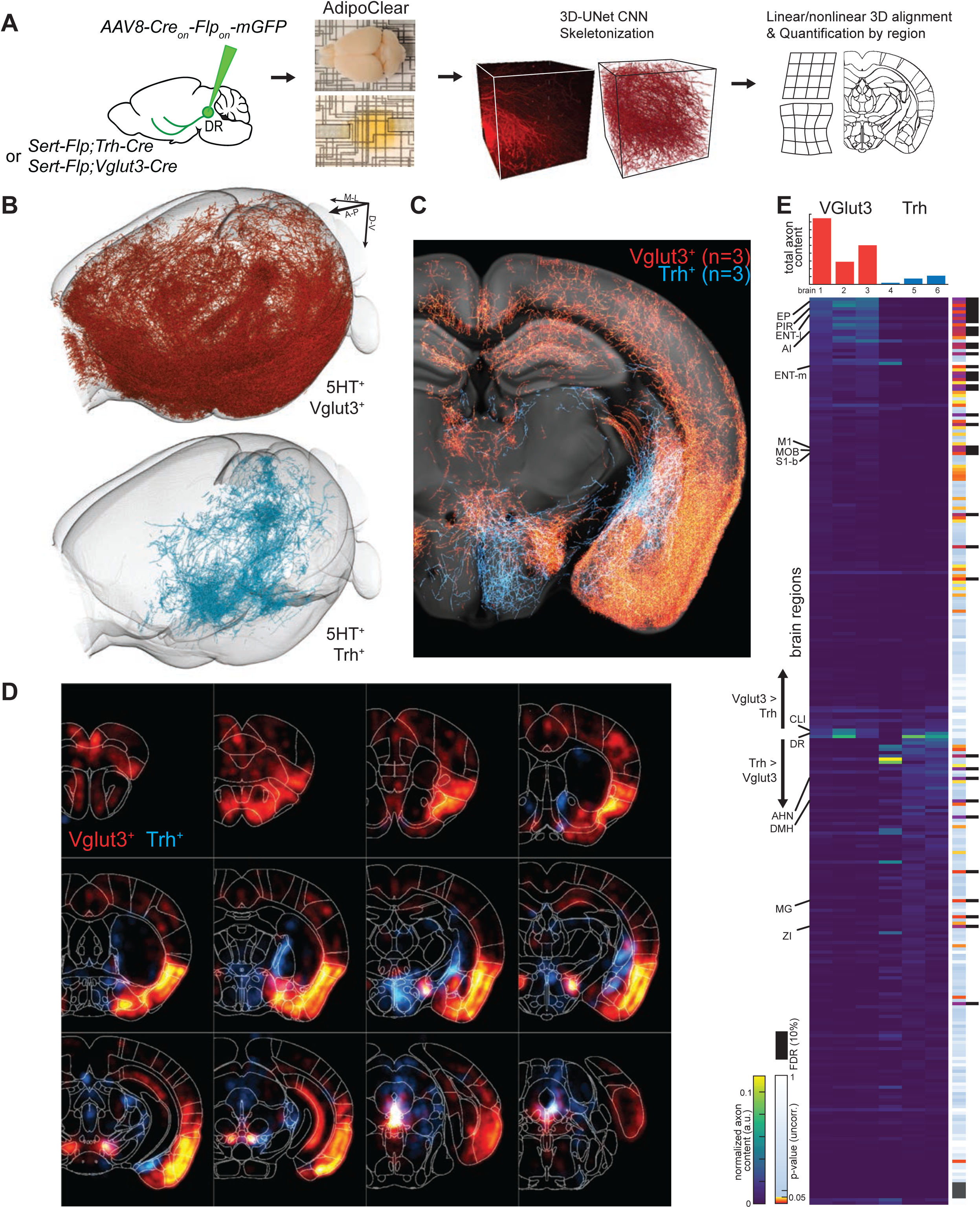
Whole-brain projectome of *Trh^+^* and *Vglut3^+^* serotonin neuron populations. **(A)** Experimental schematic outlining the intersectional viral strategy, brain clearing, automated 3D axon segmentation, and alignment to the Allen Brain Institute Common Coordinate Framework. **(B)** Axonal innervation in a 3D view of the left hemisphere of one representative brain each from the intersection of *Sert-Flp* and either *Vglut3-Cre* or *Trh-Cre*. **(C)** Coronal Z-projection (500 µm) of depth axonal innervation patterns of 6 aligned brains. The schematic reference image is one 5 µm thick plane in the middle of the 500 µm stack. **(D)** Coronal heatmaps of axonal innervation patterns at 12 positions along the rostral–caudal axis for the same 6 brains as seen in (C). Weightings for individual voxels represent axonal content within a radius of 225 µm. **(E)** Top, bar plot shows the quantification of total axonal content in each of 6 brains prior to normalization. Bottom, heatmap breaks out the total content into each of 282 individual brain regions using boundaries from the Allen Institute CCF. Values are normalized to both target region volume and total axon content per brain. Display order is grouped by mean normalized prevalence of axons in each genotype and ordered by the second principal component. P-values for individual t-tests are uncorrected; those that survive FDR-testing at 10% are indicated with a black bar. See ***Supplemental Table 4*** for full list of regions. EP, Endopiriform nucleus; M1, Primary motor area; S1-B, Primary somatosensory area, barrel field; CLI, Central linear nucleus raphe; AHN, Anterior hypothalamic nucleus; DMH, Dorsomedial nucleus of the hypothalamus; MG, Medial geniculate nucleus; ZI, Zona incerta.

We visualized the axon terminals in brain regions targeted by either the *Trh^+^* or the *Vglut3^+^* population of serotonin neurons. Initial assessment of selected target regions suggested a strong segregation of axonal projection patterns between these two populations (***Figure 7B,C***), consistent with data from tissue sections (***Figure 6D***). Axons from both populations followed similar initial trajectories and were observed in shared structures along the length of the median forebrain bundle (***Figure 7C,D***; ***Figure 7—figure supplement 1–3; Video 1***). However, as was seen in tissue sections, there were extensive differences in innervation patterns of terminal axon fields between the *Vglut3^+^* and *Trh^+^* populations. Whole-brain quantitative and statistical analyses showed *Vglut3^+^* axons preferentially in anterolateral cortical regions and adjacent structures such as olfactory bulb, agranular insular cortex, endopiriform, piriform, and claustrum as well as other cortical regions such as entorhinal, primary motor, and barrel cortices. By contrast, Trh^+^ axons were largely absent from these structures. Conversely, subcortical regions primarily in thalamus (zona incerta and medial geniculate) and hypothalamus (anterior and dorsomedial nuclei) were preferentially targeted by *Trh^+^* axons and largely avoided by the *Vglut3^+^* population (***Figure 7E; Video 1***).

Given the variability of locations and amount of viral transduction, individual brains from the same genotype exhibit considerable variation in total axons labeled (***Figure 7E*** top) and in detailed projection patterns (***Figure 7—figure supplement 1–2***). These variabilities further highlighted regions that were targeted densely but exclusively in one or two individual brains Notable examples include anterior bed nucleus of stria terminalis (BNST), posterior amygdala, and globus pallidus external segment (GPe) in *Trh^+^* projections and the lateral central amygdala (CeA) and dentate nucleus of the cerebellum for *Vglut3^+^* projections. While we did identify large-scale patterns of collateralization for these two subtypes of serotonin neurons, one possible contribution to this inter-individual variability in projection patterns is heterogeneity within molecularly defined subpopulations of serotonin neurons.

### Whole-brain axonal arborization patterns of individual serotonin neurons

Our whole-brain projection analyses indicate that axonal arborization patterns of molecularly defined serotonin neuron subpopulations are still very complex (***Figure 7***). To examine the extent to which this reflects projection patterns of individual serotonin neurons, we combined the cell-type-specific sparse labeling strategy we recently developed (Lin et al., 2018) (***Figure 8—figure supplement 1A***) and the fluorescence micro-optical sectioning tomography (fMOST) platform (Gong et al., 2016). We fully reconstructed whole-brain arborization patterns of 50 DR serotonin neurons from 19 *Sert-Cre* mice. All reconstructed neurons and their process were registered to the Allen Reference Atlas (***Figure 8A***) (Gilbert, 2018). The cell bodies of these 50 serotonin neurons covered large regions of the DR (***Figure 8—figure supplement 1B***), and their projections collectively innervated the majority of the brain regions (***Figure 8—figure supplement 1C***), suggesting their broad coverage of DR serotonin neuron types. Complete morphological reconstruction revealed that the projection pattern of individual DR serotonin neurons was highly diverse yet follow some general patterns (***Figure 8—figure supplement 1C***). We categorized them into six groups based on their projection patterns. The groups were named after the destination innervated by the highest proportion of their terminals (***Figure 8B–G***; ***Figure 8—figure supplement 2–3***; ***Video 2***; **Materials and methods**). We highlight some of features below.

**Figure 8.**
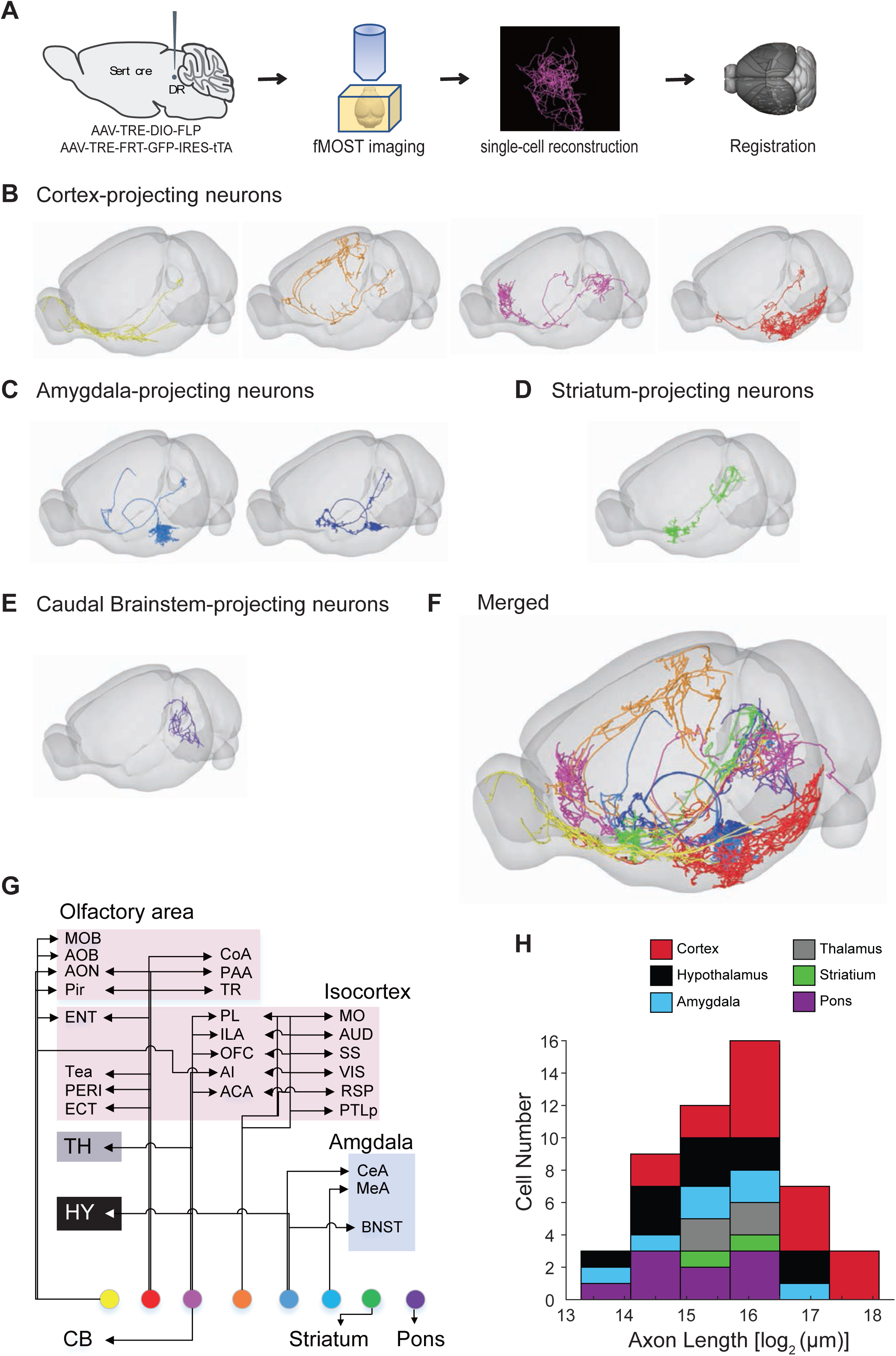
Whole-brain axonal arborization patterns of individual serotonin neurons. **(A)** Schematic of single-neuron reconstruction pipeline. **(B)** Four examples of cortex-projecting DR serotonin neurons, projecting primarily to olfactory cortex and olfactory bulb (1^st^), dorsal cortex (2^nd^), prefrontal cortex (3^rd^), and entorhinal cortex (4^th^). **(C)** Two examples of amygdala-projecting DR serotonin neurons. **(D)** A striatum-projecting DR serotonin neuron. **(E)** A caudal brainstem-projecting DR serotonin neuron. **(F)** Merged example neurons from panels B–E. **(G)** Schematic diagram illustrating the major projection targets of 8 sample neurons in panel F. (**H**) Histogram showing the distribution of cell numbers according to the total axon length.

#### Cortex-projecting

This group of DR serotonin neurons (n = 17) all dedicated more than 50% their total axonal length to innervate cortical regions. In general, they had the most complex branching pattern, with various combinations of target regions (***Figure 8B, G***; ***Figure 8—figure supplement 2A***). This group could be further divided into four subgroups: olfactory area-projecting (containing >50% of the axon length within in olfactory cortex, n = 7), prefrontal cortex-projecting (>30% of the axon length, n = 4), entorhinal cortex-projecting (>30% of the axon length, n = 3), and dorsal cortex-projecting (including motor, somatosensory, retrosplenial, and visual cortex; n = 3). In the olfactory area-projecting subgroup, three brains had branches in the olfactory bulb (OB). Axons of the three dorsal cortex-projecting neurons traveled the largest distance, entering the cortex anteriorly and extending to the posterior end. Interestingly, seven cortex-projecting serotonin neurons also sent substantial branches to non-cortical regions (>10% of axon length), including three that innervated the cerebellum.

#### Hypothalamus-projecting

Each cell of this group (n = 11) dedicated more than 33% axon length to innervate the hypothalamus (***Figure 8B***; ***Figure 8—figure supplement 2B***). None of them had collateralization in the cortical regions. Four cells had substantial branches to the pons, one to the amygdala and one to the thalamus (> 10% axon length).

#### Amygdala-projecting

The axons of all amygdala-projecting DR serotonin neurons (n = 7; ***Figure 8C***; ***Figure 8—figure supplement 3A***) followed either or both of the two distinct routes to reach the amygdala (***Figure 8—figure supplement 4A1, A2***). Six cells had collateralized axonal terminals in the hypothalamus (>10% axon length). One cell sent a branch to the contralateral cortex. Two cells had branches that were almost equally distributed in the bed nucleus of stria terminalis (BNST), central amygdala (CeA), and medial amygdala (MeA). Interestingly, the axon arbors at the CeA or MeA were highly restricted and dense (***Figure 8—figure supplement 4A3***).

#### Other groups

The rest of the DR serotonin neurons were divided into thalamus-projecting, striatum-projecting, and caudal brainstem-projecting groups based on the highest density of axons (***Figure 8—figure supplement 3B–D***), even though most also innervated other regions including the medulla and spinal cord (***Figure 8—figure supplement 1C***). While most projections of the DR serotonin neurons were unilateral (37/50), one of the thalamus-projecting neurons had symmetrical bilateral projections in the thalamic target (***Figure 8—figure supplement 4B***). For the two DR serotonin neurons that innervated ventral striatum (also called nucleus accumbens, or NAc), arborization appeared to be restricted to either the core or the shell (***Figure 8—figure supplement 4C***).

In summary, these results revealed remarkable complexity and heterogeneity of serotonin neuron projections at the single-cell level. They nevertheless followed certain patterns. For example, the cell body locations of these traced neurons (***Figure 8—figure supplement 1B***) were consistent with our previous results (Ren et al., 2018) as well as single-cell transcriptome, HCR-smFISH, and whole-brain bulk tracing data. Specifically, cortex-projecting serotonin neurons were biased towards ventral pDR (14 out of 17 cells), whereas amygdala-projecting ones were preferentially distributed in the dorsal pDR (5 out of 7 cells).

Forebrain-projecting serotonin neurons have been estimated to have very long axons based on the total serotonin fiber density and number of serotonin neurons (Wu et al., 2014). With single-cell tracing, we could now directly quantify the total length of axons of the 50 DR serotonin neurons. We found that the total axon length of these DR serotonin neurons exhibited considerable heterogeneity, from 1.2 cm to 22.7 cm. When examined across the six groups, cortex-projecting serotonin neurons indeed had the longest axons (***Figure 8H***). The longest serotonin neuron has 22.7 cm total axon length, or >10 times the length of a mouse brain in the longest dimension from rostral olfactory bulb to caudal brainstem.

## Discussion

Collectively, ∼12000 serotonin neurons from the DR and MR in mice (∼0.015% of all CNS neurons) innervate the entire forebrain (Ishimura et al., 1988) and modulate diverse physiological and behavioral functions. A fundamental question is how these serotonin neurons are organized to manage such a broad range of modulation. Single-cell transcriptomics have emerged in recent years as a powerful means to classify neuronal types, supplementing traditional criteria based on developmental history, morphology, projection patterns, and physiological properties (Luo et al., 2018). Using high-depth single-cell RNAseq in combination with systematic fluorescence in situ hybridization, whole brain projection mapping via intersectional methods and single-axon tracing, we begin to shed light on the relationship between transcriptomic clusters, the spatial location of their cell bodies, and brain-wide projection patterns of serotonin neurons.

### Relationship between molecular identity and cell body distribution

Our single-cell transcriptome analysis identified 11 molecularly distinct types of serotonin neurons in the DR and MR (***Figure 1***). Based on tissue source from which scRNA-seq data was collected and fluorescent in situ hybridization using transcriptomic cluster markers, we were able to assign six types to principal DR (pDR), one type to caudal DR (cDR), and four types to MR (***Figure 3***). The fact that we can assign specific transcriptomic clusters to specific groups of raphe nuclei indicate that molecularly defined serotonin populations are spatially segregated at least at this coarse level. Our results are broadly consistent with previous findings that utilized developmental origin to differentiate raphe serotonin neurons (Okaty et al., 2015).

The six types of serotonin neurons within principal DR exhibit further specificity in spatial distributions. Specifically, serotonin neurons from DR-1–3 clusters are preferentially localized in dorsal pDR, whereas those from DR-4–6 in ventral pDR, with DR-6 neurons preferentially localized to the ventral lateral wings. These data support and extend our previous finding (Ren et al., 2018) for the preferential ventral pDR location of *Vglut3^+^* serotonin neurons, which is highly expressed in DR-4 and DR-5 clusters. Together, these findings revealed the molecular basis for the differentiation of dorsal/ventral DR sub-systems (***Figure 2 and 3***).

cDR has been suggested to be more similar to the MR than to pDR in their connectivity (Commons, 2015; Kast et al., 2017). Our single-cell transcriptomic analysis indicated that serotonin neurons in the cDR are strikingly homogenous and profoundly different from both pDR and MR at the molecular level (***Figure 1***, ***Figure 4***, ***Figure 4—figure supplement 1***). All cDR serotonin neurons co-express *Vglut3*, and express several unique markers **(*Figure3—figure supplement 2*)**. By contrast, we could not discern obvious difference in spatial distributions among the four MR types, despite the fact that MR serotonin neurons have heterogenous developmental origin (Jensen et al., 2008). It will be interesting to examine in the future whether molecularly defined MR serotonin neurons have specific axonal projection patterns.

We note that in a recent high-quality comprehensive whole-brain scRNA-seq study, Zeisel et al. (2018) grouped 437 brainstem serotonin neurons in the dataset (presumably containing the entire B1–B9 groups) into just five clusters. Compared to droplet-based platform of Zeisel et al., our FACS-based scRNA-seq platform allowed more sequencing depth per cell (***Figure 1—supplement 1B***) and sequencing larger number of cells, accounting for our finer classification of transcriptomic types of serotonin neurons. This illustrates the value of using genetically targeted strategies to characterize important but rare types of cells in the brain.

### Relationship between molecular identity and projection-defined serotonin sub-systems

In our previous study, we characterized two projection-defined parallel DR serotonin sub-systems. We found that serotonin neurons that project to the orbitofrontal cortex (OFC) and central amygdala (CeA) differ in input and output patterns, physiological response properties, and behavioral functions (Ren et al., 2018). Whole-brain collateralization patterns of these two sub-systems indicate that there must be additional sub-systems projecting to regions not visited by either of these two sub-systems projects to, including much of the thalamus and hypothalamus. What is the relationship between molecularly defined serotonin neurons and projection-defined sub-systems?

Using viral-genetic intersectional approaches to access specifically *Vglut3^+^* pDR serotonin neurons in combination with staining in histological sections (***Figure 6***) and iDISCO-based whole brain imaging (***Figure 7***), we found that these neurons project profusely to cover much of the neocortex, as well as the olfactory bulb, cortical amygdala, and lateral hypothalamus. Comparisons of the projection patterns of *Vglut3^+^* (this study) with OFC-projecting DR serotonin neurons (Ren et al., 2018) suggest that the latter is a large subset of the former. Brain regions that are innervated by *Vglut3^+^* but not by OFC-projecting serotonin neurons include the somatosensory barrel cortex, ventral striatum, and a specific subset of CeA. Interestingly, our previous study indicated that ∼23% of CeA-projecting DR serotonin neurons are *Vglut3^+^* (Ren et al., 2018), so it is possible that even within a refined nucleus like CeA, molecularly distinct serotonin projections are confined to sub-regions with a finer resolution.

We also assessed the whole-brain projection patterns of a largely complementary population of DR serotonin neurons, namely those that express *Trh* and thus belong to DR-1–3 clusters. We found that *Trh^+^* serotonin neurons predominantly project to subcortical regions, most notably anterior and medial nuclei of the hypothalamus and several thalamic nuclei, a pattern mostly complementary to the *Vglut3^+^* population (***Figures 6–7***, ***Video S1***). Given that our previous CeA-projecting DR serotonin neurons do not innervate most of the hypothalamus, and *Trh^+^* serotonin neurons only partially innervate CeA, these two populations are at most partially overlapping.

These comparisons support a broad correspondence between molecular identity and axonal projection patterns at the level of DR serotonin neuronal populations that include multiple transcriptomic clusters. These results will enable future testing of whether a more precise correspondence exists at the level of single transcriptomic clusters that we have defined here. Our transcriptome data suggest that each DR/MR serotonin neuron type can be distinguished from others by the expression of two marker genes (***Figure 1D; Figure3—figure supplement 1–3***), supporting the view that neuronal subtypes are generally specified by unique combination of genes rather than single genes (Li et al., 2017b). With the generation of drivers based on these marker genes, intersectional methods in combination with location-specific viral targeting can be used in the future to dissect the projection patterns of the individual transcriptomic clusters.

### Insights from single-cell reconstruction

The complexity of whole-brain projection patterns of *Vglut3^+^* and *Trh^+^* populations discussed above can be driven by: 1) heterogeneity of projection patterns of different transcriptomic clusters within the *Vglut3^+^* or *Trh^+^* population; 2) heterogeneity of projection patterns of serotonin neurons within the same transcriptomic cluster; 3) complex collateralization patterns of individual serotonin neurons; and 4) a combination of some or all of the above. If scenario 1 were the only contributing factor, then there would be only six different projection patterns for the pDR serotonin neurons. However, our single-cell reconstruction of DR serotonin neurons revealed many more than 6 branching patterns (e.g., ***Figure 8G***), indicating that there must be diverse collateralization patterns even within the same transcriptomic cluster, and highlighting the complexity of individual serotonin neuron projections. Despite the complexity, these single-cell tracing data nevertheless suggest certain rules obeyed by serotonin neurons.

First, there is a general segregation of cortical- and subcortical-projecting serotonin neurons. Of the 46 forebrain-projecting DR serotonin neurons, 31 have a strong preference (>90% total axon length) for innervating either cortical or subcortical regions. This is likely an underestimate of the preference, especially for cortical-projecting ones, as their axons necessarily need to travel through the subcortical regions to reach the cortex. (As a specific example, most forebrain-projecting DR serotonin neurons pass through the lateral hypothalamus to reach their targets; it is thus difficult to determine whether axons in the lateral hypothalamus play a local function.) Second, most of the serotonin neurons tend to focus a majority of their arborization within one brain region (e.g., prefrontal vs entorhinal cortex, ***Figure 8B*;** CeA vs MeA, ***Figure 8—figure supplement 4A3***). The subcortical-projecting serotonin neurons appear to have more specificity, with most neurons exhibiting dense arborization in one or two nuclei. The cortical-projecting serotonin neurons can have elaborate arborization patterns across multiple cortical areas (e.g., ***Figure 8—figure supplement 1C***) and the longest axon lengths per cell (***Figure 8H***).

Together with our study on the projection-defined serotonin sub-systems (Ren et al., 2018), we believe it is unlikely that the major function of the forebrain-projecting serotonin system is to broadcast information non-discriminately. Our study is limited by the scope (50 reconstructed cells out of 9000 DR serotonin neurons). To fully reveal the organizational logic of the serotonin system, efforts should be put into larger scale single-cell reconstruction integrated with molecular identity and functional studies of individual transcriptomic clusters of serotonin neurons.

### Integrating multiple features within individual serotonin sub-systems

The molecular features of different serotonin cell types suggest their functional properties. For example, several studies have reported that subgroups of serotonin neurons in the MR and DR express Vglut3 and indeed, subsequent slice recording confirmed that serotonin terminals can co-release glutamate and serotonin. In addition to neurochemical properties, each serotonin neuron population reveals a specific combination of distinct genes responsible for electrophysiological (ion channels), connectivity (wiring molecules), and functional (neurotransmitter/peptide receptors) properties (***Figure 4*; *Figure 4—figure supplement 1A***). For example, most *Crhr2^+^* neurons co-express *Vglut3* and *Npy2r*, which suggests that these serotonin neurons use glutamate as co-transmitter in their cortical targets, and are in turn modulated by corticotropin-releasing hormone and neuropeptide Y. Meanwhile, most *Trh^+^* serotonin neurons co-expression *Gad1*, *Kcns1*, and α1A adrenergic receptors (*Adra1a*) specifically. We can speculate that these serotonin neurons use Trh (and perhaps GABA) as co-transmitters to regulate the physiology of their subcortical targets, and are in turn modulated by locus coeruleus norepinephrine neurons.

Our previous study suggests that DR serotonin sub-systems have biased input but segregated output (Ren et al., 2018). Here we found that each of the transcriptomic clusters of serotonin neurons have distinct combination of axon guidance and cell adhesion molecules **(*Figure 4—figure supplement 1A*)**. These differentially expressed wiring molecules might be used during development to set up distinct projection patterns of different serotonin neuron types (Deneris and Gaspar, 2018; Kiyasova and Gaspar, 2011; Maddaloni et al., 2017), and/or used in adults to maintain wiring connectivity or contribute to the remarkable ability of serotonergic axons to regenerate after injury (Jin et al., 2016).

In conclusion, our comprehensive transcriptomic dataset and its 11 distinct groups of the DR and MR serotonin neurons reveals the molecular heterogeneity of the forebrain-projecting serotonin system. We have shown that the molecular features of these distinct serotonin groups reflect their anatomical organization and provide tools for future exploration of the full projection map of molecularly defined serotonin groups. The molecular architecture of serotonin system lays the foundation to integrate connectivity, neurochemical, physiological, and behavioral functions. This integrated understanding of serotonin can in turn provide novel therapeutic strategies to treat brain disorders involving this important neuromodulator.

## Materials and methods

### Animals

All procedures followed animal care and biosafety guidelines approved by Stanford University’s Administrative Panel on Laboratory Animal Care and Administrative Panel of Biosafety in accordance with NIH guidelines. For scRNA-seq (Figure 1 and 4), male and female mice aged 40–45 days on a C57BL/6J background were used. The *Ai14* tdTomato Cre reporter mice (JAX Strain 7914) and *Sert-Cre* (MMRRC, Stock #017260-UCD) were crossed and the offspring was used where indicated. For HCR experiments (Figures 2) wild-type male and female mice aged 8 weeks on a C57BL/6J background were used. For whole brain axon tracing experiments (Figures 5–7), male and female mice aged 8–16 weeks on a CD1 and C57BL/6J mixed background were used. The *Vglut3-Cre* (also known as *Slc18a8-Cre*; JAX Strain 18147), *Thr-ires-Cr*e (JAX Strain 032468; gift from Dr. Bradford B. Lowell), *Sert-Flp* (generated in this study), *H11-CAG-FRT-stop-FRT-EGFP* (generated in this study), *IS* (JAX Strain 028582; gift from Dr. Z. Josh Huang) were used where indicated. For single-cell reconstruction experiments, male and female mice aged 6–16 weeks on a CD1 and C57BL/6J mixed background were used. Mice were group-housed in plastic cages with disposable bedding on a 12 hours light/dark cycle with food and water available *ad libitum*.

#### Generation of *Sert-Flp* mice

*Sert-Flp* was generated by the Gene Targeting and Transgenics core at Janelia Research Campus. It was generated by homologous recombination in embryonic stem cells using standard procedures. A cassette of *IRES-FlpO-loxP2272-ACE-Cre POII NeoR-loxp2272* was inserted after the TAA stop codon of *Sert.* Targeting was verified in embryonic stem cells by long-arm PCR. After microinjection, chimaeras were bred with CD-1 females and F1 offspring were screened by long-arm PCR to identify mice with germline transmission of the correctly targeted construct.

#### Generation of H11-CAG-FRT-stop-FRT-EGFP mice

*H11-CAG-FRT-stop-FRT-EGFP* was generated using site-specific integrase-mediated transgenesis via pronuclear injection (Tasic et al., 2011) by Stanford transgenic facility. The *CAG-FRT-stop-FRT-EGFP* cassette was inserted as a single-copy transgene at the *H11* locus (Tasic et al., 2011).

### Transcriptome analysis

#### Single-cell isolation and sequencing

Lysis plates were prepared by dispensing 4 μl lysis buffer as described in (Tabula Muris et al., 2018). After dissociation, single tdTomato^+^ cells were sorted in 96-well plates using SH800S (Sony). Immediately after sorting, plates were sealed with a pre-labelled aluminum seal, centrifuged, and flash frozen on dry ice. cDNA synthesis and library preparation were performed using the Smart-seq2 protocol (Picelli et al., 2014). Wells of each library plate were pooled using a Mosquito liquid handler (TTP Labtech). Pooling was followed by two purifications using 0.7x AMPure beads (Fisher, A63881). Library quality was assessed using capillary electrophoresis on a Fragment Analyzer (AATI), and libraries were quantified by qPCR (Kapa Biosystems, KK4923) on a CFX96 Touch Real-Time PCR Detection System (Bio-Rad). Libraries were sequenced on the NextSeq 500 Sequencing System (Illumina) using 2 × 75-bp paired-end reads and 2 × 8-bp index reads.

#### Data processing

Sequences from the NextSeq were de-multiplexed using bcl2fastq version 2.19.0.316. Reads were aligned to the mm10 genome using STAR version 2.6.1a (Dobin et al., 2013). Gene counts were produced using HTseq version 0.11.2 (Anders et al., 2015) with default parameters, except ‘stranded’ was set to ‘false’, and ‘mode’ was set to ‘intersection-nonempty’. Genes located on Y chromosome were removed from the count table to exclude sex bias.

#### Clustering

Standard procedures for filtering, variable gene selection, dimensionality reduction and clustering were performed using the Seurat package version 3.0 (Butler et al., 2018; Stuart et al., 2019). Specifically, cells with fewer than 300 detected genes were excluded. A gene was counted as expressed if it has at least one read mapping to it and is detected in at least 3 cells. Cells with fewer than 50,000 reads were excluded. Counts were log-normalized for each cell using the natural logarithm of 1 + counts per million [ln(CPM+1)]. All genes were projected onto a low-dimensional subspace using principal component analysis. Cells were clustered using a variant of the Louvain method that includes a resolution parameter in the modularity function (Tabula Muris et al., 2018) and visualized using a 2-dimensional t-distributed Stochastic Neighbour Embedding (tSNE) of the PC-projected data. Molecularly distinct cell populations were assigned to each cluster based on differentially expressed genes. Plots showing the expression of the markers for each cell subtype appear in the ***Figure 3—figure supplement 1–3*.**

#### Gene co-expression networks

The relationship between gene expression was measured using rank correlation statistics. Pearson correlations were computed across all cells. We first removed low expressed genes by selecting genes with mean expression CPM>2, leaving ∼8000 genes in the dataset. Pearson correlation coefficients (*rp*) were computed for each gene and significance was tested by bootstrapping (1,000 iterations). A correlation table containing *rp* above 0.3 and below –0.3 can be found in ***Supplemental Table 3***. Reported values are mean from the bootstrapped values. Gene functional categories were retrieved from HGNC resource (https://www.genenames.org). Genes assigned to more than one functional category were re-assigned a single category in the following priority order: SNAREs, Secreted Ligands, ACh and Monoamine Receptors, Glutamate Receptors, Axon Guidance and Cell Adhesion Molecules (CAMs), GABA Receptors, GPCRs, Ion Channel Proteins, Transcription Factors, a full list of genes assigned to each category can be found in ***Supplemental Table 2***. Gene pairs for which *rp<0.4* were removed and remaining pairs were visualized as a network using *igraph* and *visNetwork* R packages. To further refine the final list of co-expressed genes and generate Figure 4D we focused on gene pairs for which:1) *rp>0.5*; 2) at least one gene of the pair is found among cluster markers; 3) both genes of the correlating pair belong to one of the above listed functional categories.

#### Data availability

The datasets generated and analyzed in the study are available in the NCBI Gene Expression Omnibus (GEO) (currently waiting for the access number).

## Abbreviations for anatomical regions

ACA: anterior cingulate area
AHN: anterior hypothalamic nucleus
AI: anterior insular cortex
AOB: accessory olfactory bulb
AON: The anterior olfactory nucleus
AUD: auditory cortex
BLA: basolateral amygdala
BNST: bed nucleus of the stria terminalis
BST: nucleus of stria terminalis
CB: cerebellum
cBS: caudal brainstem
CeA: central amygdala
CLI: central linear nucleus raphe
CoA: cortical amygdala
DB: nucleus of the diagonal band
DCN: deep cerebellum nuclei
dLGN: dorsal lateral geniculate nucleus
DMH: dorsomedial nucleus of the hypothalamus
DP: dorsal peduncular cortex
ECT: ectorhinal cortex
ENT: entorhinal cortex
EP: endopiriform nucleus
GU: gustatory area
HY: Hypothalamus
ILA: Infralimbic area
LHb: lateral habenula
LHy: lateral hypothalamus
LS: lateral septal nucleus
M1: primary motor area
Mbmot: midbrain, motor related
Mbsen: midbrain, sensory related
Mbsta: behavioral state related
MeA: medial amygdala
MG: medial geniculate nucleus
MOB: main olfactory bulb
NLOT: nucleus of lateral olfactory tract
NST: nucleus of solitary tract
OB: olfactory bulb
OFC: orbitofrontal cortex
ORB: orbital area
PAA: piriform– amygdalar area
PERI: perirhinal area
PERI: perirhinal cortex
PIR: piriform cortex
PL: Prelimbic area
PSTh: parasubthalamic nucleus
PTLP: Posterior parietal association area
PVH: paraventricular hypothalamus
PVHd: paraventricular hypothalamus, descending division.
RSP: retrosplenial area
S1-B: primary somatosensory area, barrel field
SI: substantia innominata
SNc: substantia nigra compacta
SNr: substantia nigra pars reticulata
SS: somatosensory
Sth: subthalamic nucleus
STR: straitum
TEA: temporal association
TH: Thalamus
TR: piriform transition area
TT: tenia tecta
VIS: visual cortex
VLPO: ventrolateral preoptic nucleus
ZI: zona incerta

## Stereotaxic Surgeries

Mice were anesthetized either with 1.5%–2.0% isoflurane and placed in a stereotaxic apparatus (Kopf Instruments). For virus injection in to the DR, the following coordinates (in mm) were used: –4.3AP, 1.10 ML, –2.85 DV; –4.3AP, 1.10 ML, –2.70 DV, with 20° ML angle. (AP is relative to bregma; DV is relative to the brain surface when AP is –1.0). After surgery, mice recovered on a heated pad until ambulatory and then returned to their homecage.

## Viral constructs

The full design of Ef1a-CreON/FlpON-mGFP is *Ef1a-fDIO-*[*membrane tag*]*-Kozak-loxP-STOPx3-loxP-ΔGFP*. The AAV vector backbones that contained the *Ef1a-fDIO were derived from pAAV-Ef1a-fDIO-EYFP* (Addgene, 27437) (Fenno et al., 2014). The [*membrane tag*]*-Kozak-loxP* sequence was synthesized by GenScript. The sequence of *loxP-ΔGFP* were cloned by PCR. And then these two pieces together with *STOPx3* were ligated into the *AAV-Ef1a-fDIO* backbone in the antisense orientation by Gibson assembly. DNA oligonucleotides were synthesized by Elim Biopharmaceuticals Inc and GenScript.

## Viruses packaging

For whole brain tracing, the AAV vector carrying *Ef1a-CreON/FlpON-mGFP* were packaged into AAV2/8 serotype with 1 x 10^12^ gc/ml by Gene Vector and Virus Core, Stanford University. 500 nl of *AAV-CreON/FlpON-mGFP* was injected into the DR for each mouse. For single cell reconstruction, AAV vectors carrying the *TRE-DIO-FlpO*, *TRE-fDIO-GFP-IRES-tTA* construct were packaged into AAV2/9 serotype with titres 10^9^–10^10^ gc/ml in Dr. Minmin Luo’s lab as described before (Lin et al., 2018). *AAV-TRE-DIO-FLPo* (10^7^ gc/ml) virus and *AAV-TRE-fDIO-GFP-IRES-tTA* (10^9^ gc/ml) virus were mixed with the ratio of 1:9. 200nL mixed virus was injected into the DR for each mouse.

## Histology and Imaging

Animals were perfused transcardially with phosphate buffered saline (PBS) followed by 4% paraformaldehyde (PFA). Brains were dissected, post-fixed in 4% PFA for 12–24 hours in 4 °C, then placed in 30% sucrose for 24–48 hours. They were then embedded in Optimum Cutting Temperature (OCT, Tissue Tek) and stored at–80°C until sectioning. For the antibody staining in Figure 1, 50-µm sections containing DR were collected onto Superfrost Plus slides to maintain the anterior to posterior sequence. All working solutions listed below included 0.2% NaN3 to prevent microbial growth. Slides were then washed 3×10 min in PBS and pretreated overnight with 0.5 mM SDS at 37°C. Slides were then blocked for 4 hours at room temperature in 10% normal donkey serum (NDS) in PBS with 0.3% Triton-X100 (PBST), followed by incubation in primary antibody (Novus, rabbit anti-Tph2) diluted 1:1000 in 5% NDS in PBST for 24 hours at RT. After 3×10 min washes in PBS, secondary antibody was applied for 6 hours at room temperature (donkey anti-rabbit, Alexa-647 or Alexa-488, Jackson ImmunoResearch), followed by 3×10 min washes in PBST. Slides were then stained for NeuroTrace Blue (NTB, Invitrogen). For NTB staining, slides were washed 1×5 min in PBS, 2×10 min in PBST, incubated for 2–3 hours at room temperature in (1:500) NTB in PBST, washed 1×20 min with PBST, and 1×5 min with PBS. Sections were additionally stained with DAPI (1:10,000 of 5 mg/mL, Sigma-Aldrich) in PBS for 10–15 min and washed once more with PBS. Slides were mounted and coversliped with Fluorogel (Electron Microscopy Sciences). After that, the slides were then imaged either using a Zeiss 780 confocal microscope or a 3i spinning disk confocal microscope (CSU-W1 SoRa), and images were processed using NIH ImageJ software. After that, whole slides were then imaged with a 5x objective using a Leica Ariol slide scanner with the SL200 slide loader.

For DR-containing slices in Figure 5 and 6, staining was applied to floating sections. Primary antibodies (Novus, rabbit anti-Tph2, 1:1000; Rockland, rabbit anti-RFP, 1:1000; Abcam, goat anti-Tph2, 1:500; Aves Labs Inc., chicken anti-GFP, 1:2000) were applied for 48 hours and secondary antibodies for 12 hours at 4°C. For serotonin terminal staining in Figure 6, floating sections were stained with Primary antibodies (Aves Labs Inc., chicken anti-GFP, 1:2000) for 60 hours and secondary antibodies for 18 hours at 4°C.

## Whole Brain Imaging of *Vglut3^+^* and *Trh^+^* Serotonin Projections

After 8-10 weeks of virus expression, mice were transcardially perfused with 20 ml 1x PBS containing 10 µg/ml heparin followed by 20 ml of 4% PFA before removing each brain and allowing them to postfix overnight at 4° C. The clearing protocol largely follows the steps outlined in (Chi et al., 2018). Brains were washed at room temperature with motion at least 30 minutes each step: 3x in 1x PBS before switching to 0.1% Triton X-100 with 0.3 M glycine (B1N buffer) and being stepwise (20/40/60/80%) dehydrated into 100% methanol. Delipidation was carried out by an overnight incubation in 2:1 mixture of DCM:methanol and a 1 hour incubation in 100% DCM the following day. After 3x washes in 100% methanol, brains were bleached for 4 hours in a 5:1 mix of methanol: 30% H2O2 and then stepwise (80/60/40/20%) restored into B1N buffer. Samples were permeabilized with 2 washes of PTxwH buffer containing 5% DMSO and 0.3M glycine for 3 hours before being washed in PTxwH overnight. Antibody labeling was carried out in PTxwH buffer at 37° C with motion. Primary antibody (Aves 1020; chicken anti-GFP, 1:1000) was added and samples were incubated for 11 days. After 3 days of washes, secondary antibody (Thermo A-31573, AlexaFluor 647 donkey anti-chicken, 1:1000) was added for 8 days, followed by another 3 days of washes. One final day of washing in 1x PBS preceded clearing. Samples were again dehydrated stepwise into methanol, using water as the alternative fraction. Delipidation proceeded as before with a DCM/methanol mixture overnight and 2x 1 hour DCM-only incubations the next day. Brains were finally cleared for 4 hours in dibenzyl ether and then stored in a fresh tube of dibenzyl ether at least 24 hours before imaging.

Samples were imaged with a LaVision Ultramicroscope II lightsheet using a 2x objective and 3 µm z-step size. Antibody fluorescense was collected from a 640 nm laser and autofluorescense captured from 488 nm illumination. Image volumes were processed and analyzed with custom Python and MATLAB scripts. In short, we trained a 3D U-Net convolutional neural network (Çiçek, et al., 2016) to identify axons in volumes and post-processed the resulting probability-based volumes as previously described (Ren et al., 2018). Using the autofluorescent channel, we aligned samples to the Allen Institute’s Common Coordinate Framework (Renier et al., 2016), applied the same transformation vectors to the volumetric projection of axons, and quantified total axon content in each brain region listed in ***Supplemental Table 4***.

## Hybridization Chain Reaction in situs

Probes were generated for use with HCR v3.0 (Molecular Technologies) (Choi et al., 2018). Wildtype 8 week old mice were perfused and brains were removed and fixed as described above. Following an overnight postfix, brains were cryoprotected in 30% sucrose until they sank and subsequently frozen at −80° C. The midbrain raphe nuclei were sectioned coronally at either 16 or 20 µm directly onto a glass slide, and dried at room temperature for 4-6 hours before storing at −20° C overnight. Sections were re-fixed again in 4% PFA, washed in 1x PBS, treated with a solution containing 50ml H2O, 0.5ml 1M Tris ph7.4, 100ul 0.5M EDTA, 35ul ProK for 6 minutes at 37° C before another round of PFA fixation and PBS washing. Hybridization and amplification steps were carried out as recommended by Molecular Technologies. Imaging was performed with a Zeiss 780 LSM confocal using a 20x objective. Double labeled cells were counted manually using the CellCounter plugin for FIJI software.

## Single Cell Reconstruction

### fMOST imaging and image preprocessing

4–6 weeks after virus injection, brains were dissected and post-fixed in 4% paraformaldehyde for 24 hr at 4 °C. The brains were rinsed in 0.01 M PBS (Sigma-Aldrich) three times (for 2 h each) and embedded in Lowicryl HM20 resin (Electron Microscopy Sciences, 14340). We use a fluorescence micro-optical sectioning tomography (fMOST) system to acquire the brain-wide image dataset at high resolutions (0.23 × 0.23×1 μm for 10 brains and 0.35 × 0.35×1 μm for the other 9 brains). Embedded brain samples were mounted on a 3D translation stage in a water bath with propidium iodide (PI). The fMOST system automatically performs the coronal sectioning with 1um steps and imaging with 2 color channels in 16-bit depth. The green channel of GFP is for visualization of neurons and the red channel of PI counterstaining is for visualization the whole brain cytoarchitecture.

### Image annotation and skeletonization

Amira software (v 5.4.1, Visage imaging, Inc) were used for semi-automatically reconstruction of single neurons. First, we use Amira to load a relatively large volume but low-resolution data and find the position of soma or major axon as a start position. Then, one by one, we load each small volume (800×800×400 voxels) of highest resolution data along the axons and dendrites to label the full structure of each neuron. We have totally reconstructed 50 high quality neurons with complete and clear axon terminals. The reconstructed neurons were checked by one another person.

### Image registration and visualization

Reconstructed neurons were aligned to the Allen Common Coordinate Framework (CCF). Elastix (Klein 2010) were used to register the manually segmented outline of a downsampled version of the PI-staining vloume (25 x 25 x 25 μm voxel size) to the outline of CCF (25 x 25 x 25μm voxel size). An initial affine registration was applied for align the sample to CCF, followed by an iterative non-rigid b-spline transformation for more precise registration. The generated parameters were used for coordinate transformation of reconstructed neurons to align them to the CCF. Cinema 4d (MAXON Computer GmbH, Germany) and customized python scripts were used to generate 3D rendered images and movies.

### Data analysis and quantification

We use customized program in MATLAB (MathWorks) for the fiber lengths calculation and heatmap generation. The projection diagram was drawn with Microsoft Office Visio.

## Acknowledgements

We thank Caiying Guo and the Howard Hughes Medical Institute/Janelia Research Campus for assisting the generation of the *Sert-Flp* mice, Yanru Chen and the Stanford transgenic core for assisting the generation of the *H11-CAG-FRT-stop-FRT-EGFP* mice, Bradford B. Lowell for *Thr-ires-Cre* mice, Z. Josh Huang for the *IS* mice, Stanford Gene Vector and Virus Core for producing virus, Kang Shen and Nathan McDonald for sharing and advising the 3i spinning disc confocal microscope, Yanwen Sun for assisting the generation of a customized program related to 3D images processing, Tie-Mei Li for assisting the cloning of *Ef1a-CreON/FlpON-mGFP*.

## Additional information

### Author contributions

J.R. and A.I. collected single cell samples. A.I. and S.S.K. processed the cells for single-cell RNA-seq, and A.I. performed scRNA-seq data analysis, with support from S.R.Q. D.F. performed FISH experiments and analyzed data, with assistance from S.G. J.R. developed and characterized *Sert-Flp* mice and intersectional viral-genetic strategies. D.F. and J.R. performed whole-brain intersectional axon tracing experiments with assistance from S.G. D.F. analyzed whole-brain axon maps using algorithms originally developed by A.P. G.Q.Z. generated the Flp reporter mouse with the support from J.L.R. J.Z. performed the bulk of single axon tracing, with assistance from R.W. and R.L. and support from M.L. P.L. and A. L. performed fMOST imaging with the support from Q.L. L.L. supervised the project. J.R. and L.L. wrote the manuscript with contribution from all authors especially A.I., D.F., J. Z., and S.R.Q.

---

**Figure 1—figure supplement 1.**
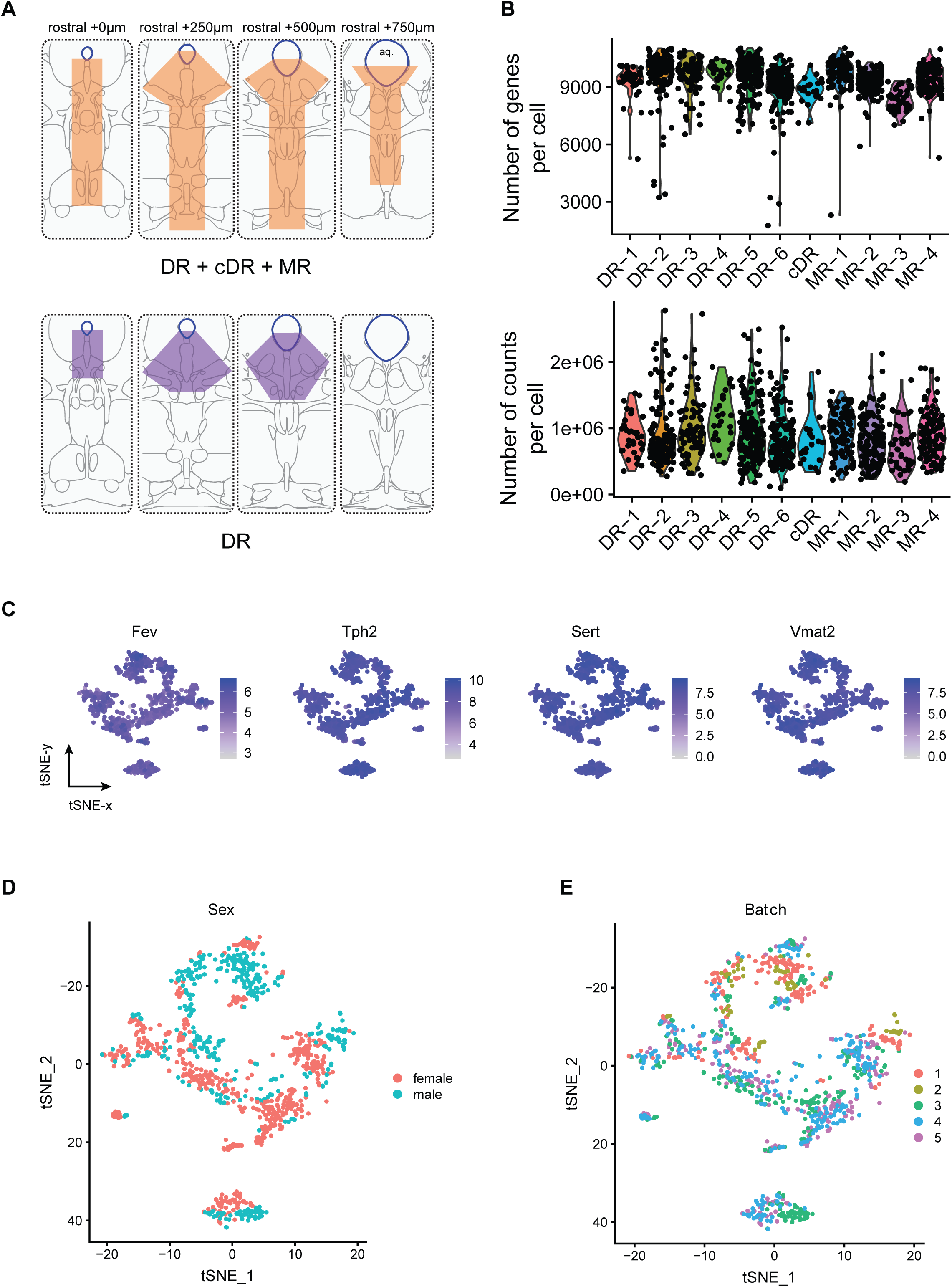
Technical characteristics of scRNA-seq experiments. **(A)** Schematic illustration of the sampling strategy. Up, orange shadows indicate dissected regions in the coronal brainstem sections in the first set of experiments. These regions contained the entire MR and DR. Bottom, purple shadows indicate dissected regions in the coronal brainstem sections in the second set of experiments. These regions only the principle DR. aq, aqueduct. **(B)** Number of genes per cell and number of reads per cells mapping to exons across 11 clusters. (**C)** Expression of *Fev*, *Tph2*, *Sert* and *Vmat2* across individual cells. Cells are colored by log-normalized expression of each transcript, and the color legend reflects the expression values of ln(CPM+1). **(D,E)** tSNE plots of 999 analyzed cells colored by sex (D) and batch (E).

**Figure 1—figure supplement 2.**
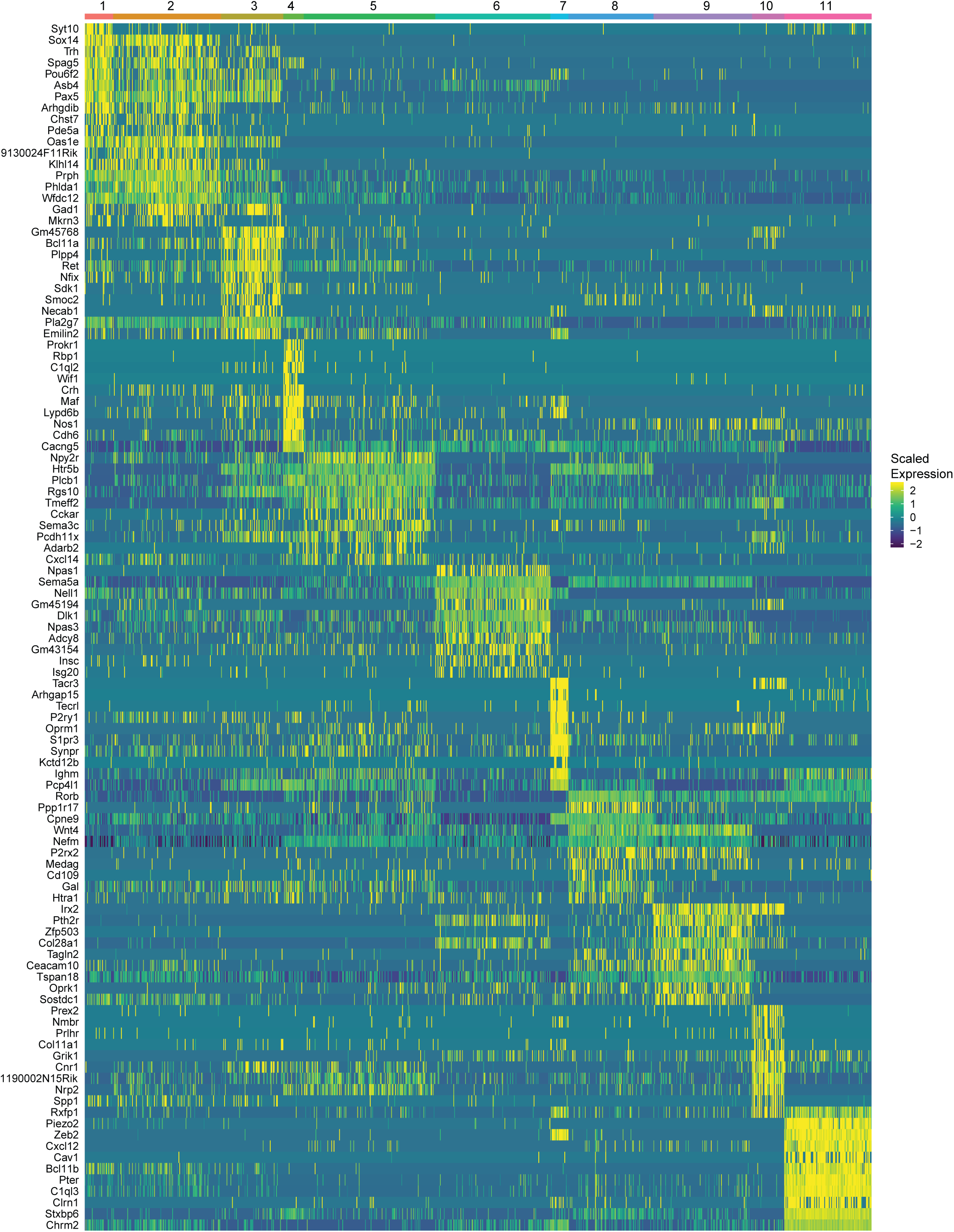
Heatmap of gene expression of top ten marker genes identified for each cluster. Expression values represent z-scores of ln(CPM+1).

**Figure 1—figure supplement 3–6.**
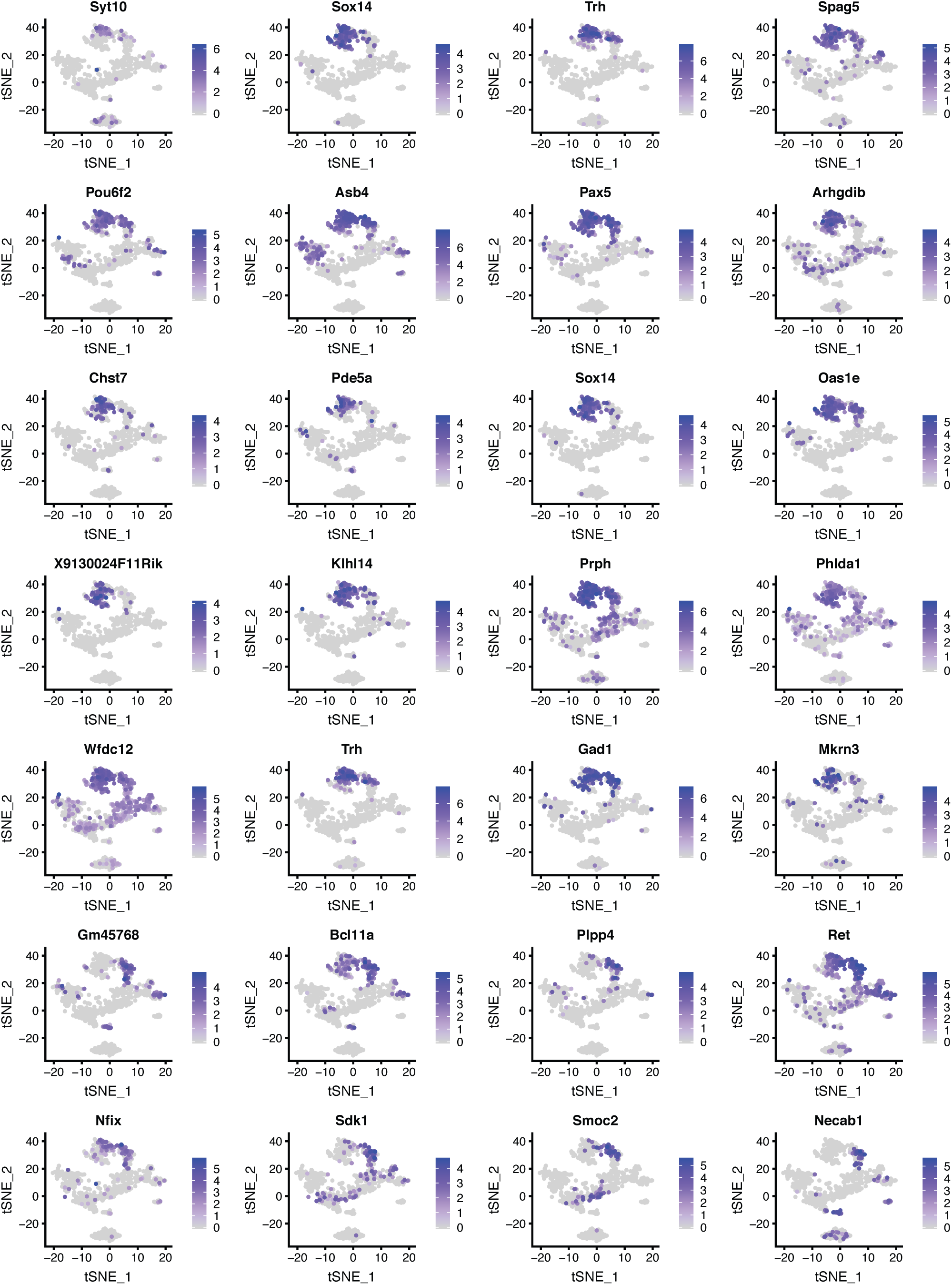
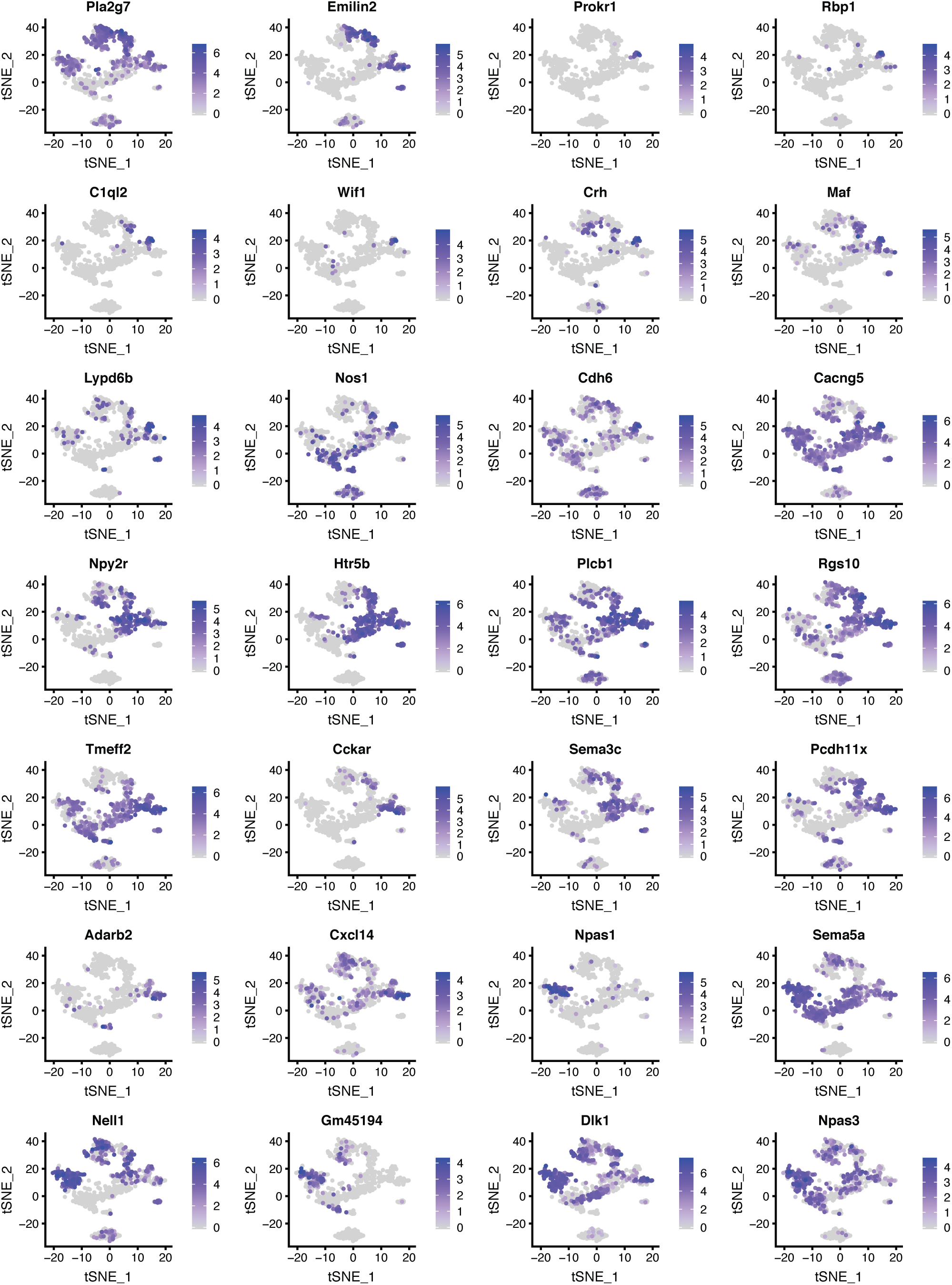
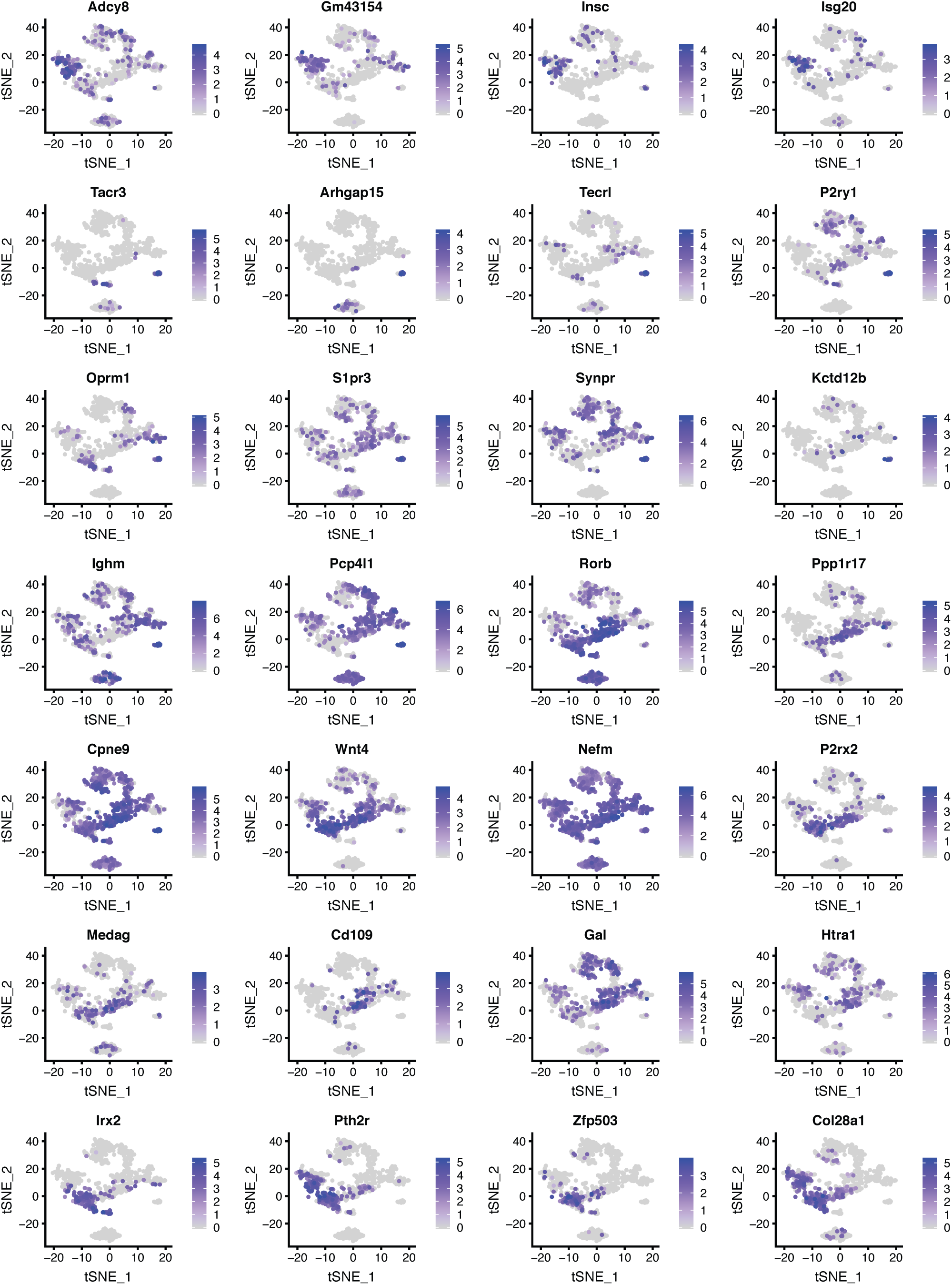
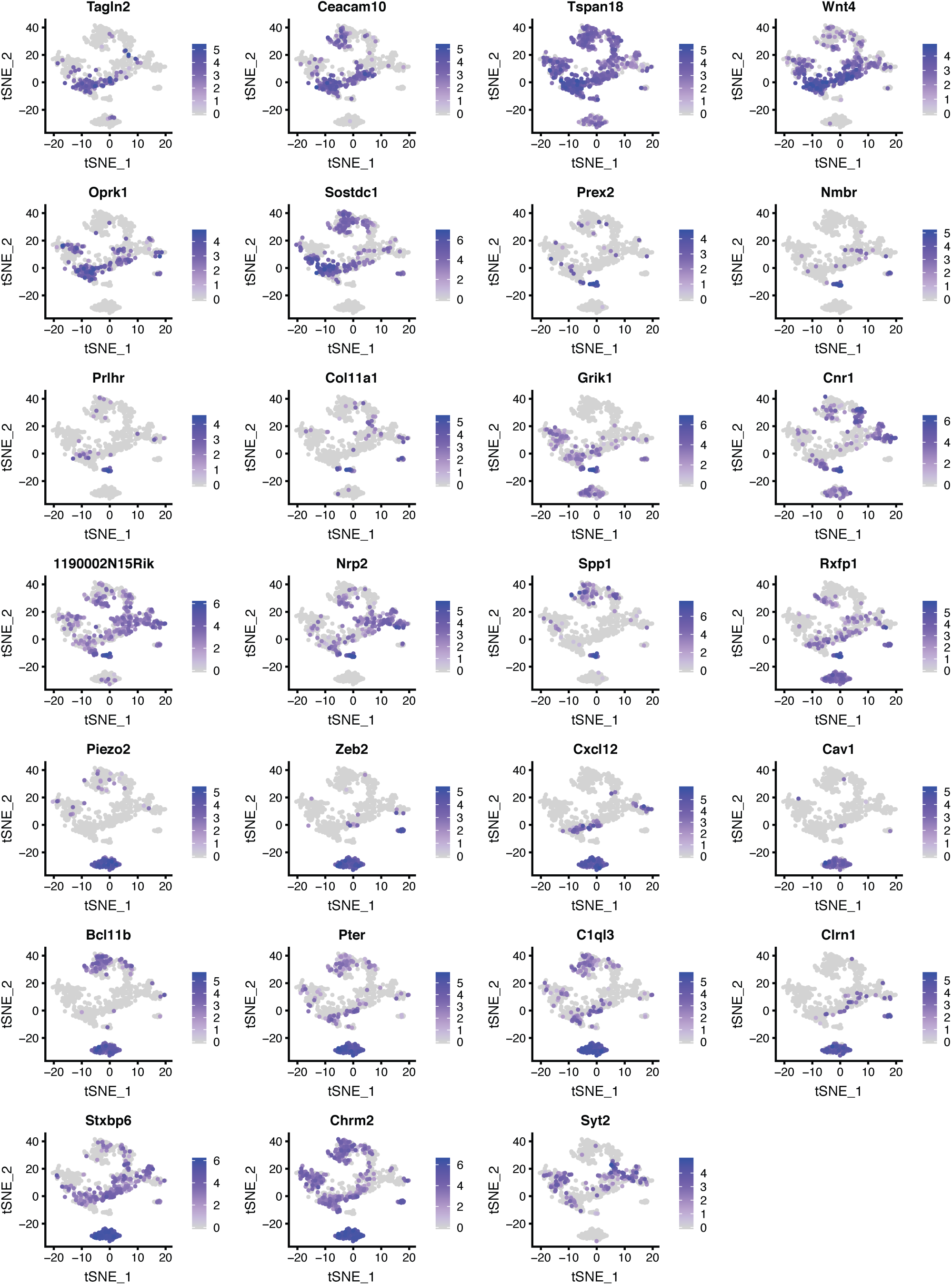
Expression patterns of selected genes in DR and MR across individual serotonin neurons presented as tSNE plots. Cells are colored by log-normalized expression of each transcript, and the color legend reflects the expression values of ln(CPM+1).

**Figure 2—figure supplement 1&2.**
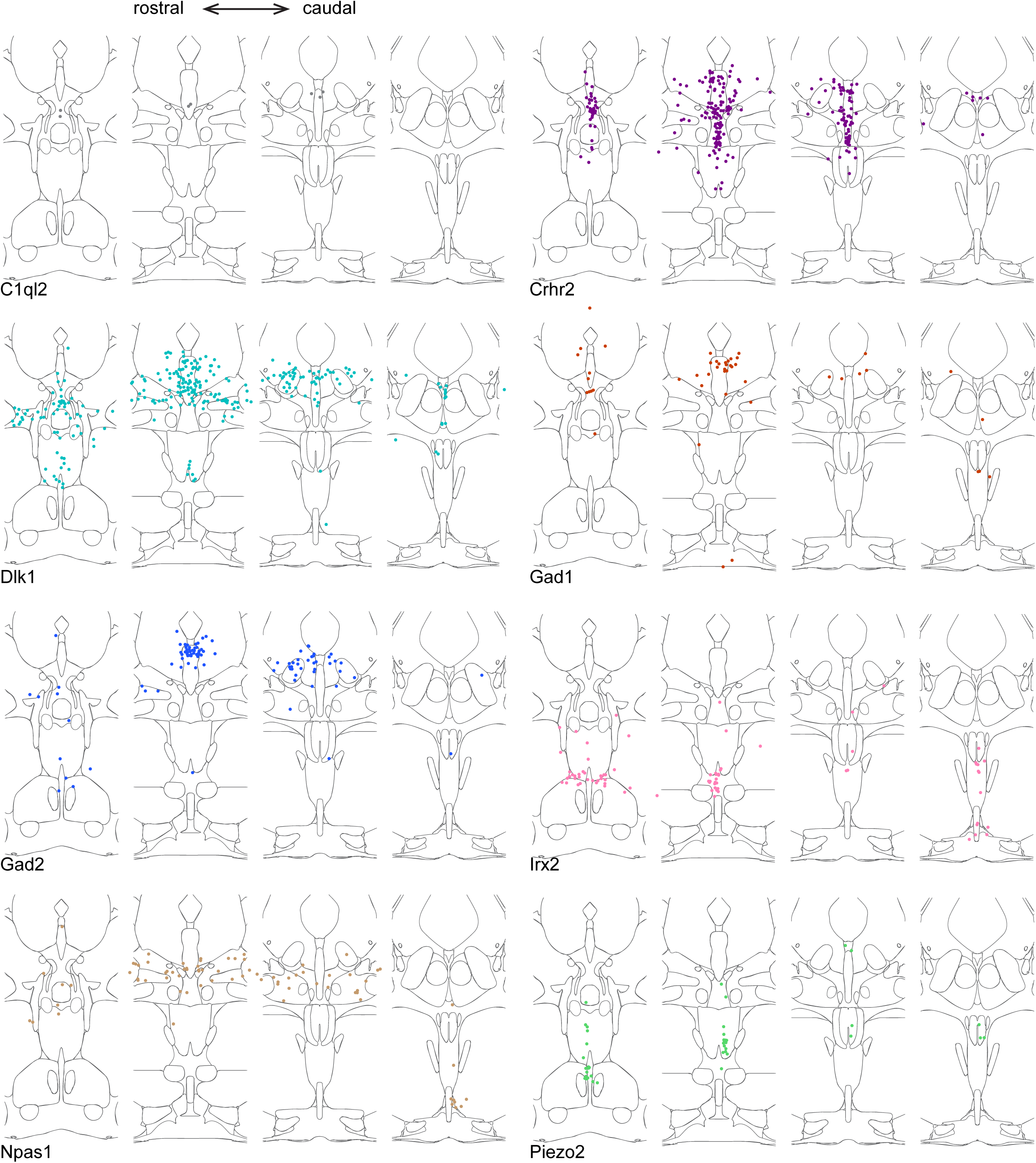
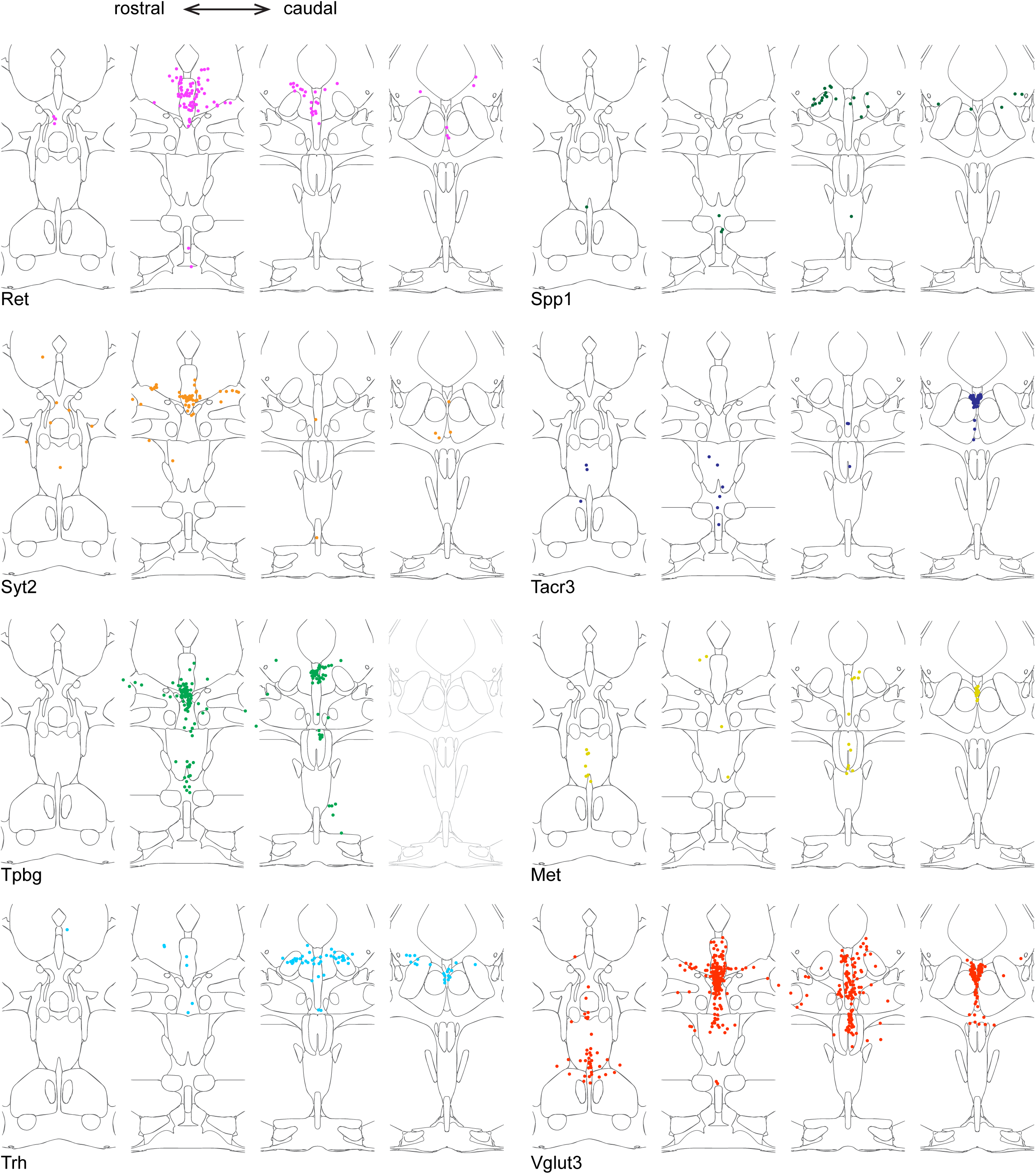
Location of cluster marker genes determined by HCR-smFISH. Schematics as described in Figure 2, broken out to show double-positive cells for Tph2 and one marker gene at a time.

**Figure 3—figure supplement 1–3.**
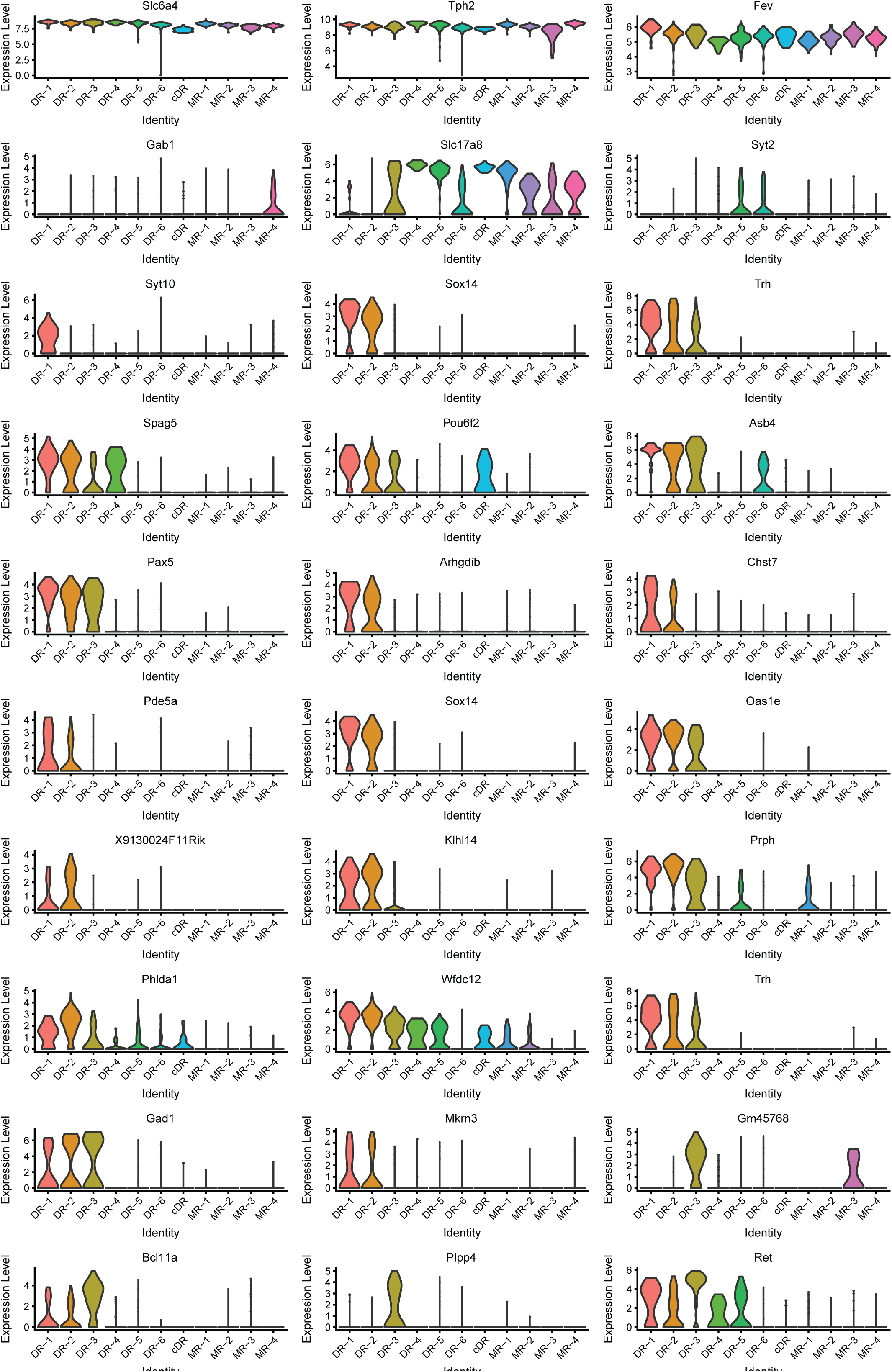
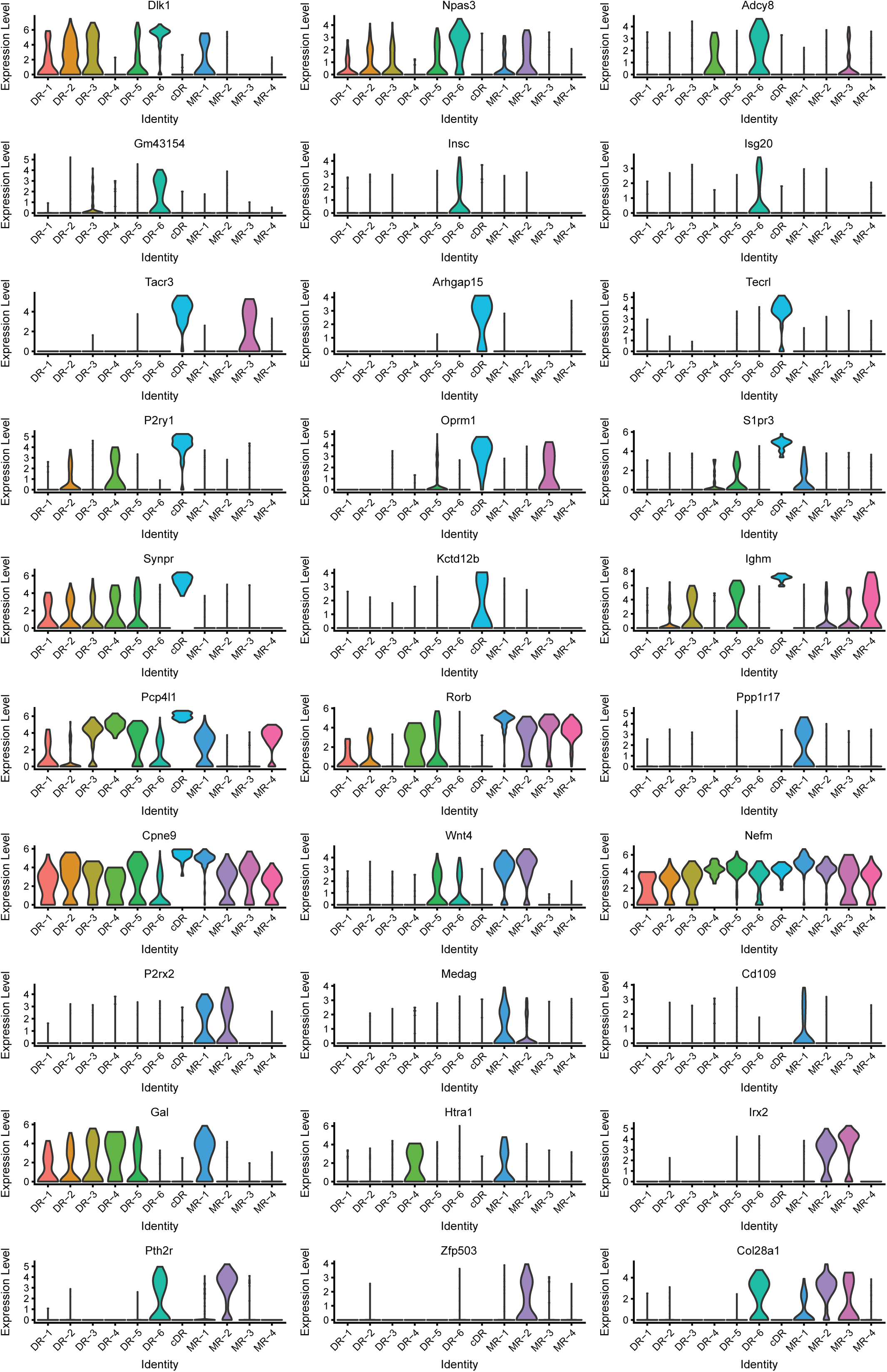
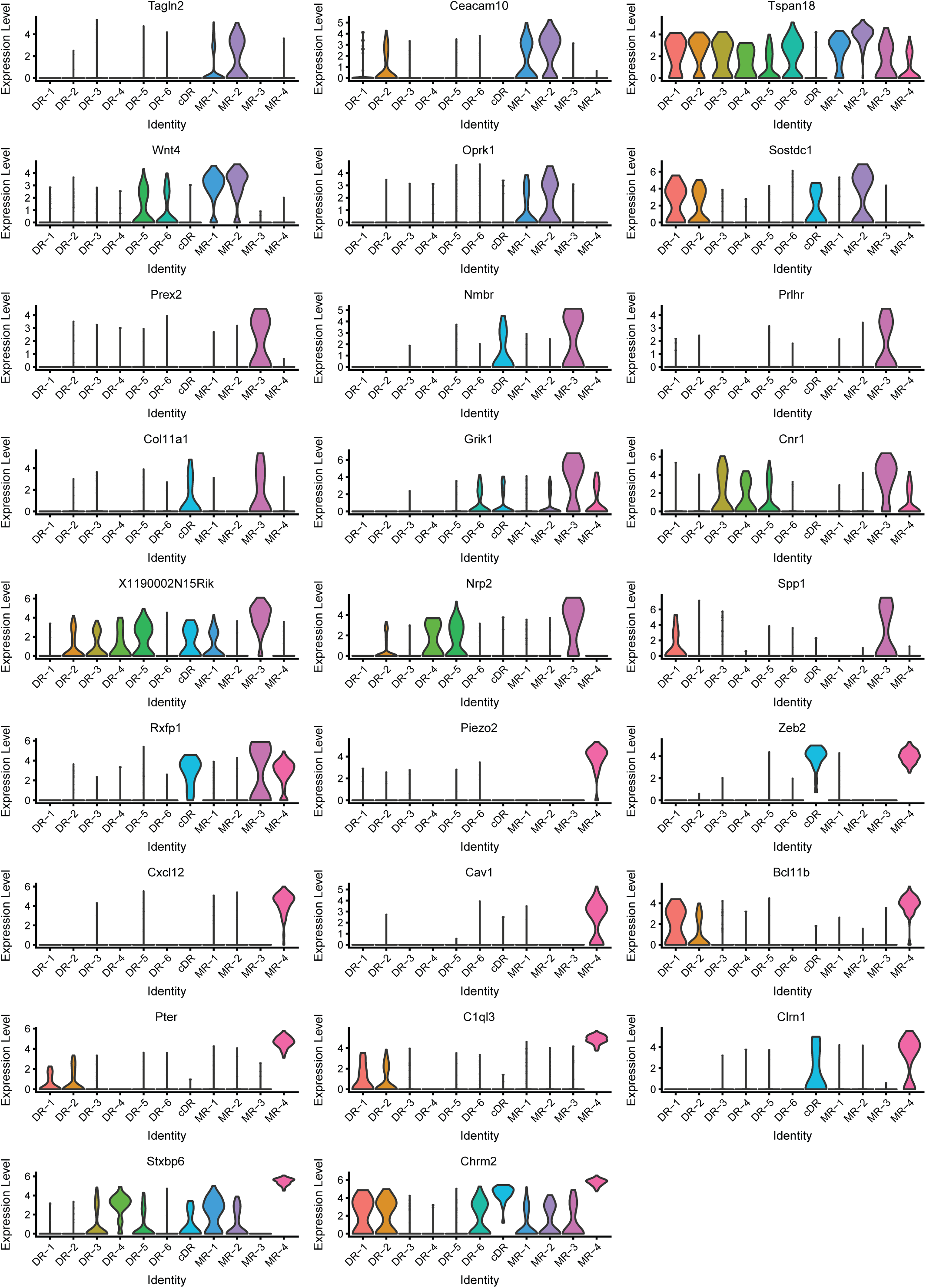
Expression of marker genes across 11 clusters. Expression levels denote for log-transformed expression [ln(counts per million+1)] of each transcript.

**Figure 4—figure supplement 1.**
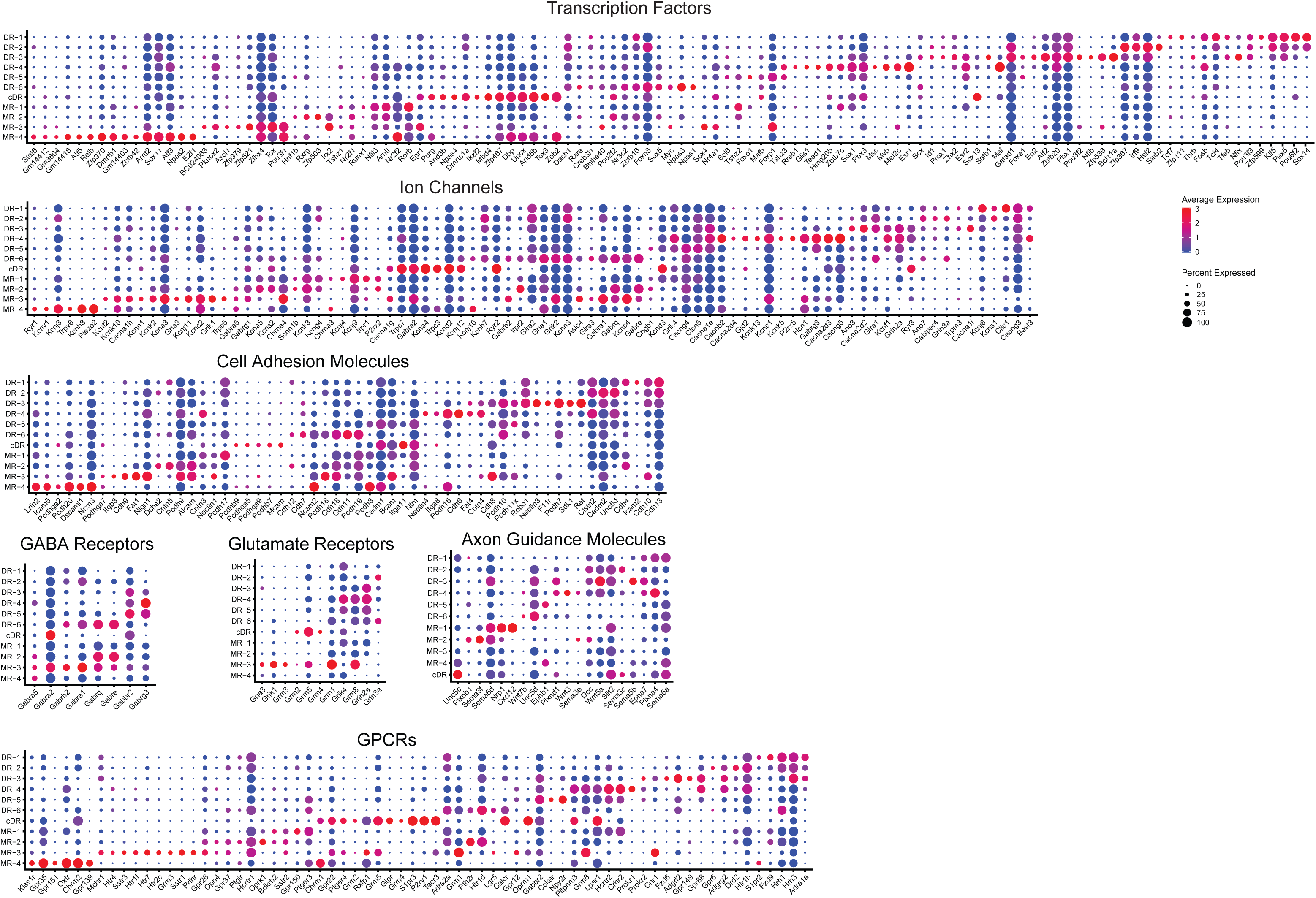
**Expression of the most variable genes among the listed functional categories across 11 molecularly distinct Tph2+ clusters.**

**Figure 4—figure supplement 2.**
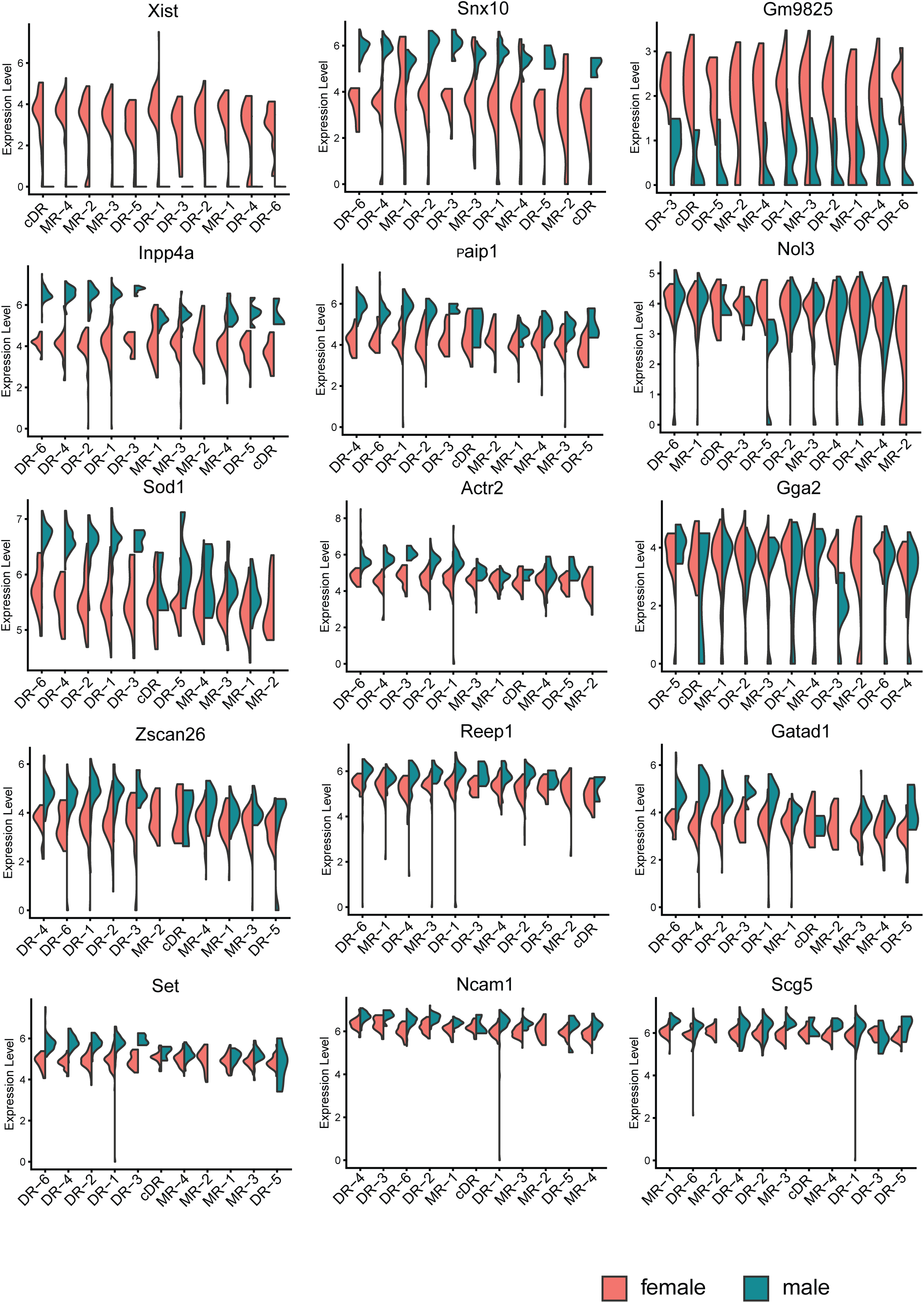
Sexually dimorphic genes consistently detected across the majority of serotonin cell subtypes. Expression levels denote for log-transformed expression [ln(CPM+1)] of each transcript.

**Figure 7—figure supplement 1.**
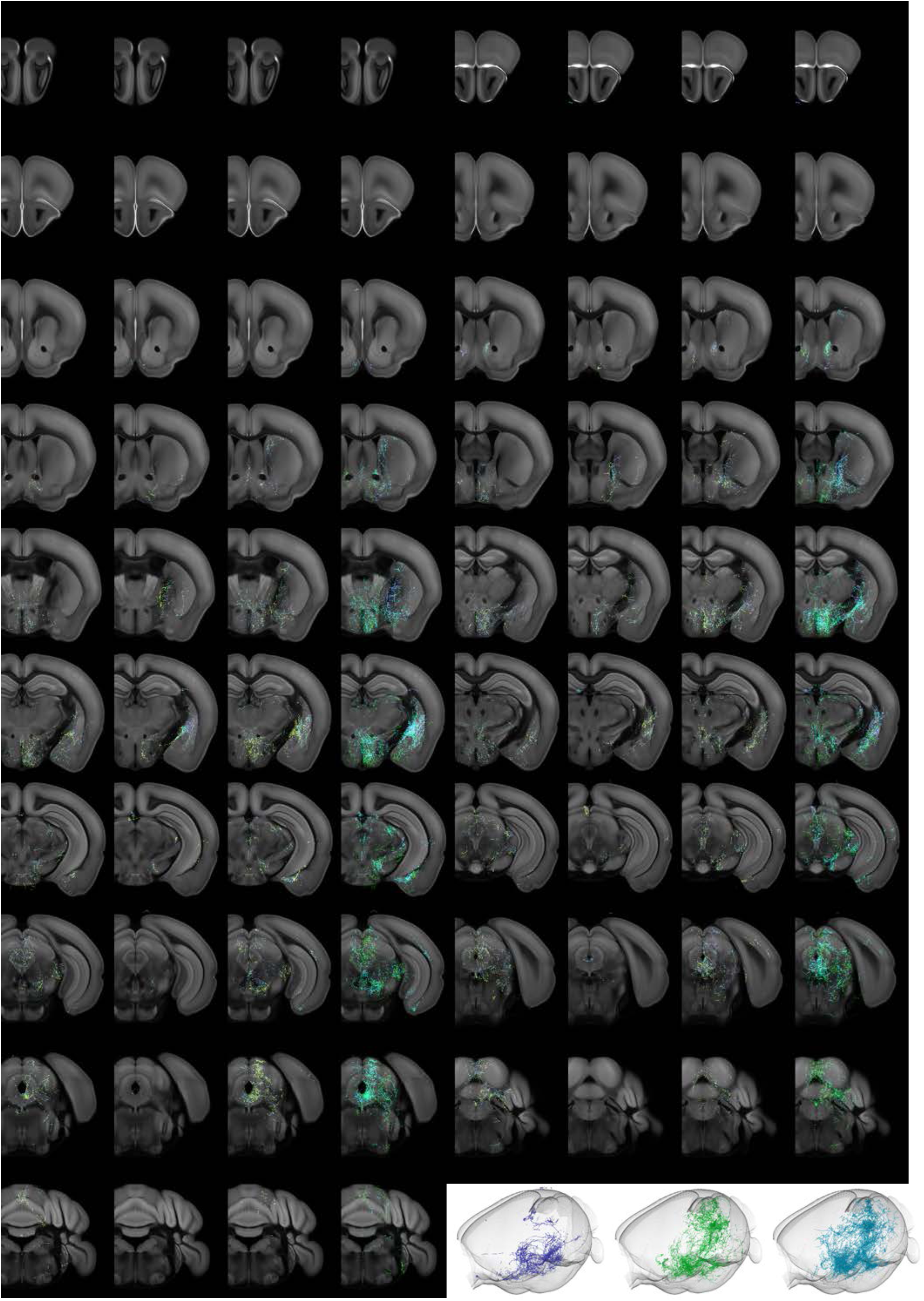
Individual projectome variability of the Trh+ serotonin population. Coronal slices of data along the rostral–caudal axis, at spacing of 500 µm. The three left panels are 500 µm Z-projections of individual brains, color-coded by Z-depth. Merge image shows the same volume of the three slices overlaid and colored individually by brain. Lower right, 3D view of axonal content for each of the three brains.

**Figure 7—figure supplement 2.**
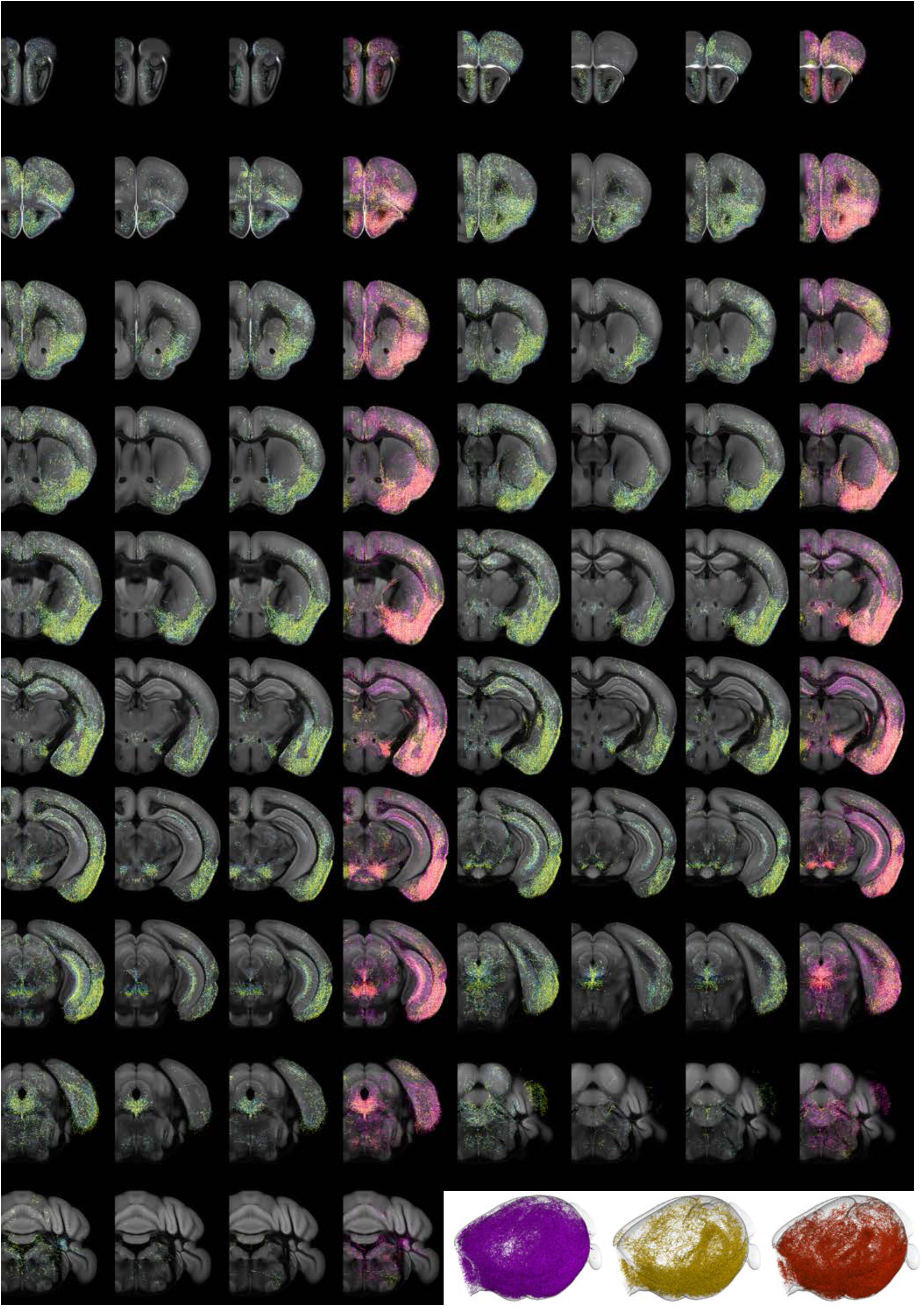
Individual projectome variability of the Vglut3+ serotonin population. Coronal slices of data along the rostral–caudal axis, at spacing of 500 µm. The three left panels are 500 µm Z-projections of individual brains, color-coded by Z-depth. Merge image shows the same volume of the three slices overlaid and colored individually by brain. Lower right, 3D view of axonal content for each of the three brains.

**Figure 7—figure supplement 3.**
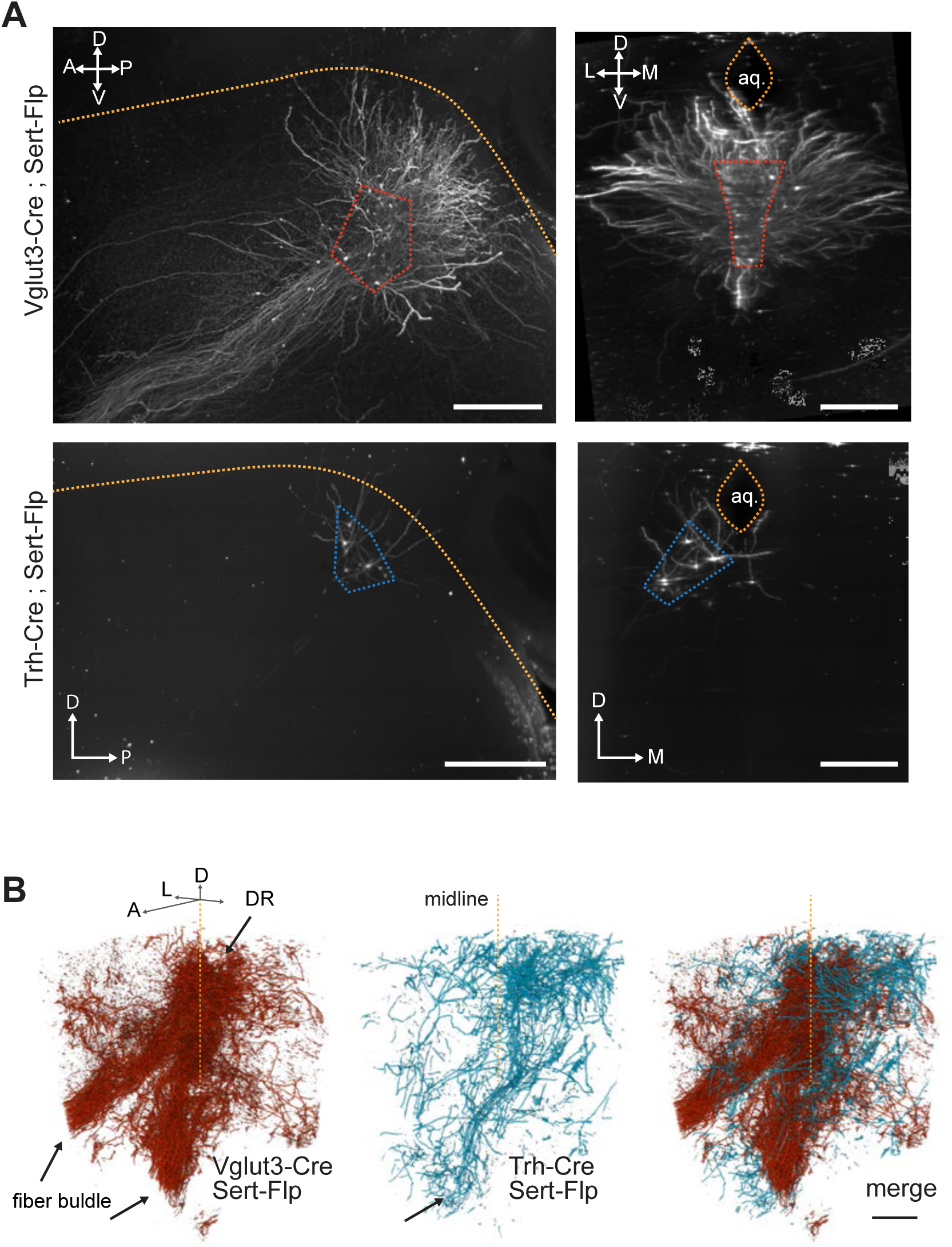
The cell body location and initial axonal segments of the Trh+ and Vglut3+ serotonin neuron subpopulations. **(A)** Z projections of cell bodies located at the injection site in DR for representative brains used in iDISCO-based whole brain imaging of axons. Dashed lines mark the ventral boundary of the aqueduct in the sagittal view and the ventral (*Vglut3-Cre*) and lateral wing (*Trh-Cre*) locations of labeled *Sert-Flp* neurons. Scale bars 500 µm. **(B)** 3D volumes of DR and initial axon branches projecting anterior and ventral into the median forebrain bundle. Volumes are from one brain each representing either *Vglut3-Cre* or *Trh-Cre* labeled *Sert-Flp* neurons. Merge at right highlights overlap. Axis labels: A, anterior; P, posterior, D, dorsal; V, ventral; M, medial; L, lateral.

**Figure 8—figure supplement 1.**
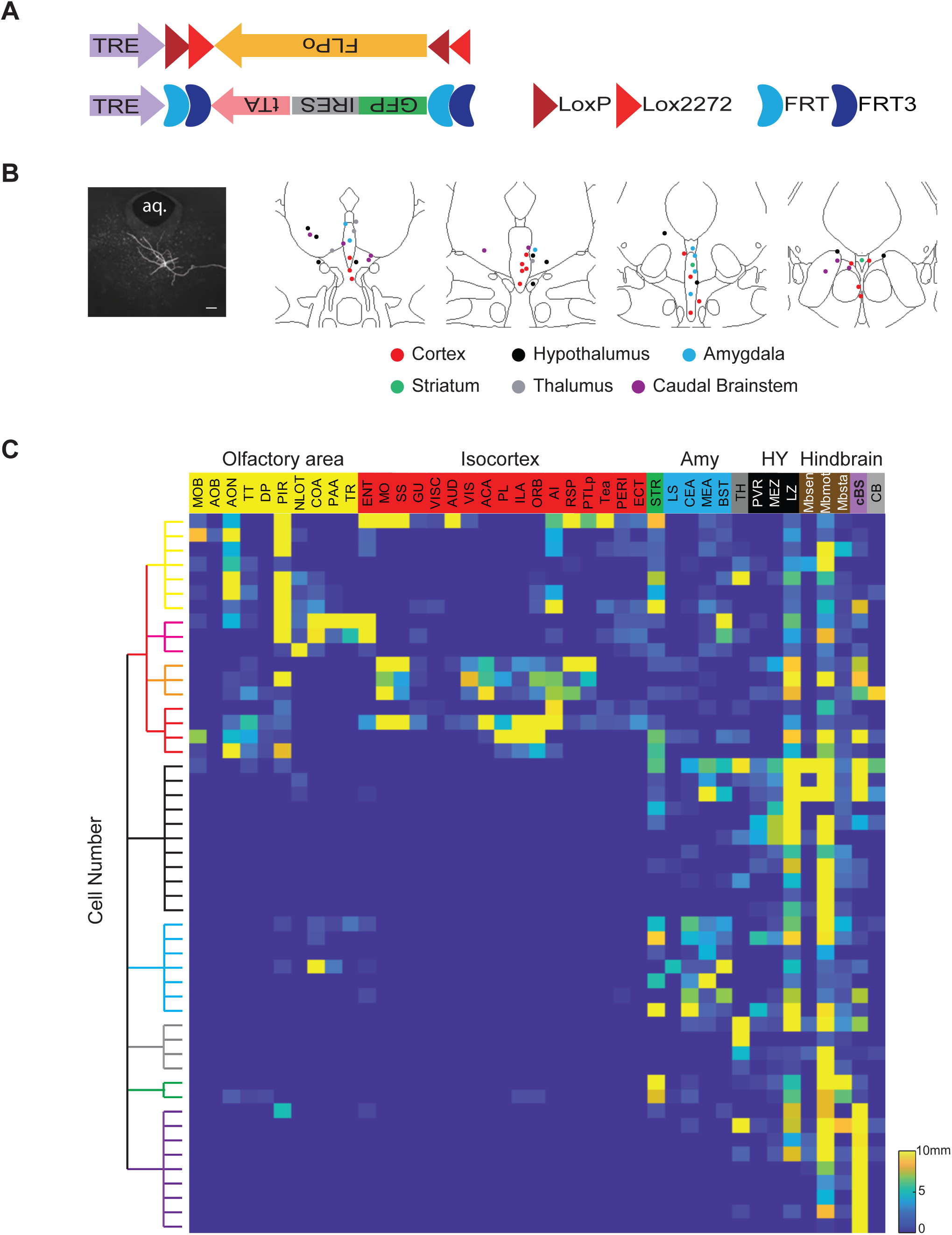
Single-cell reconstruction of DR serotonin neurons. **(A)** Design of dual-AAV sparse labeling system. **(B)** Left, an example neuron labeled by the sparse labeling system. Right, summary of the cell body position of 50 reconstructed neurons color-coded by their projection types. **(C)** Whole-brain quantification of the axon process length of the 50 reconstructed DR serotonin neurons. The 50 brains are ordered (from top to bottom) according to their projection patterns described in the text, and are the same as the numbers in Figure 8***—figure supplement 2–3***.

**Figure 8—figure supplement 2.**
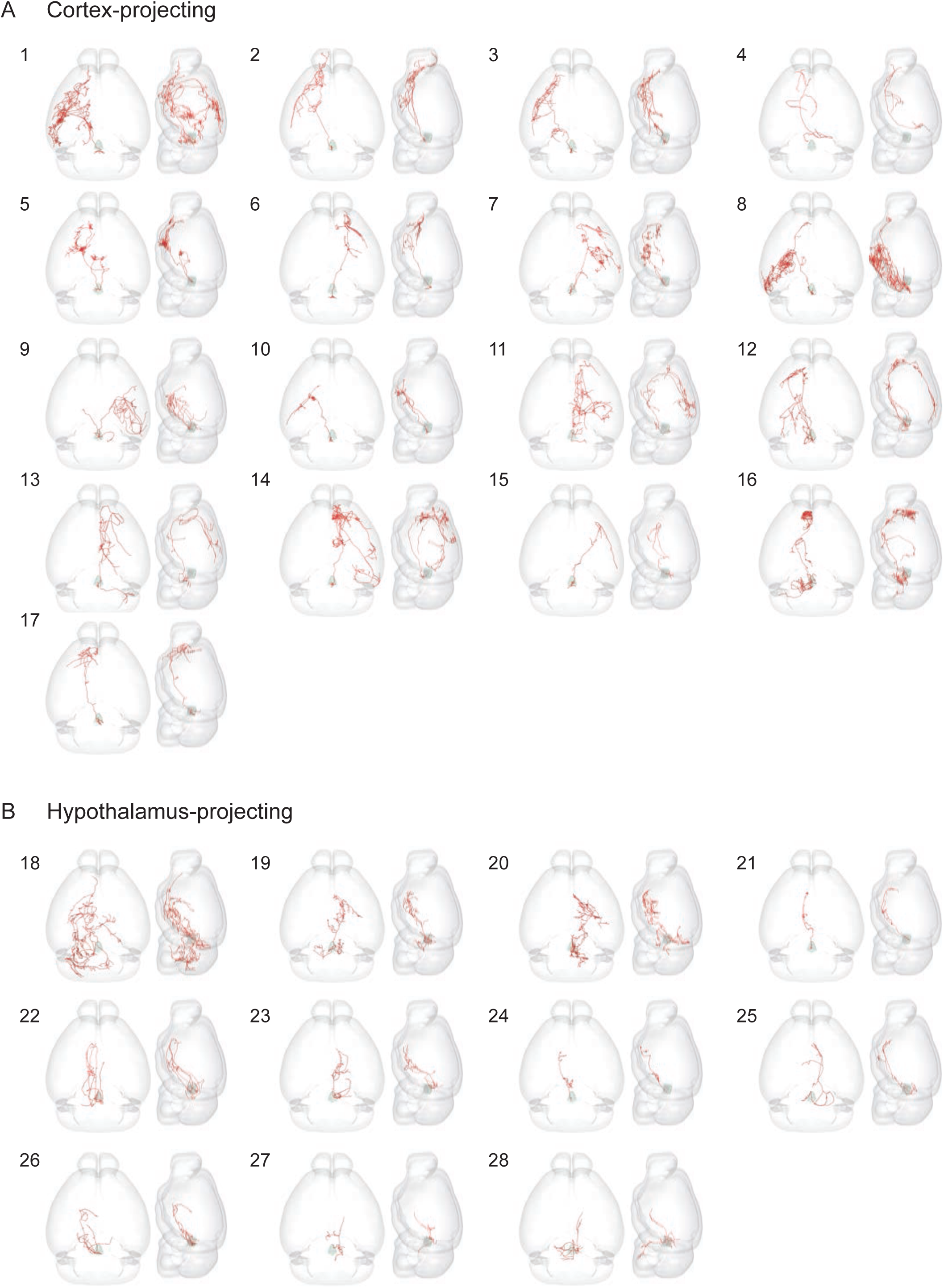
Individually reconstructed DR serotonin neurons (part I). **(A)** 17 cortex-projection DR serotonin neurons. **(B)** 11 hypothalamus-projection DR serotonin neurons. For each neuron, the left image is a horizonal view, whereas the right image is a sagittal view (ventral to the left, dorsal to the right).

**Figure 8—figure supplement 3.**
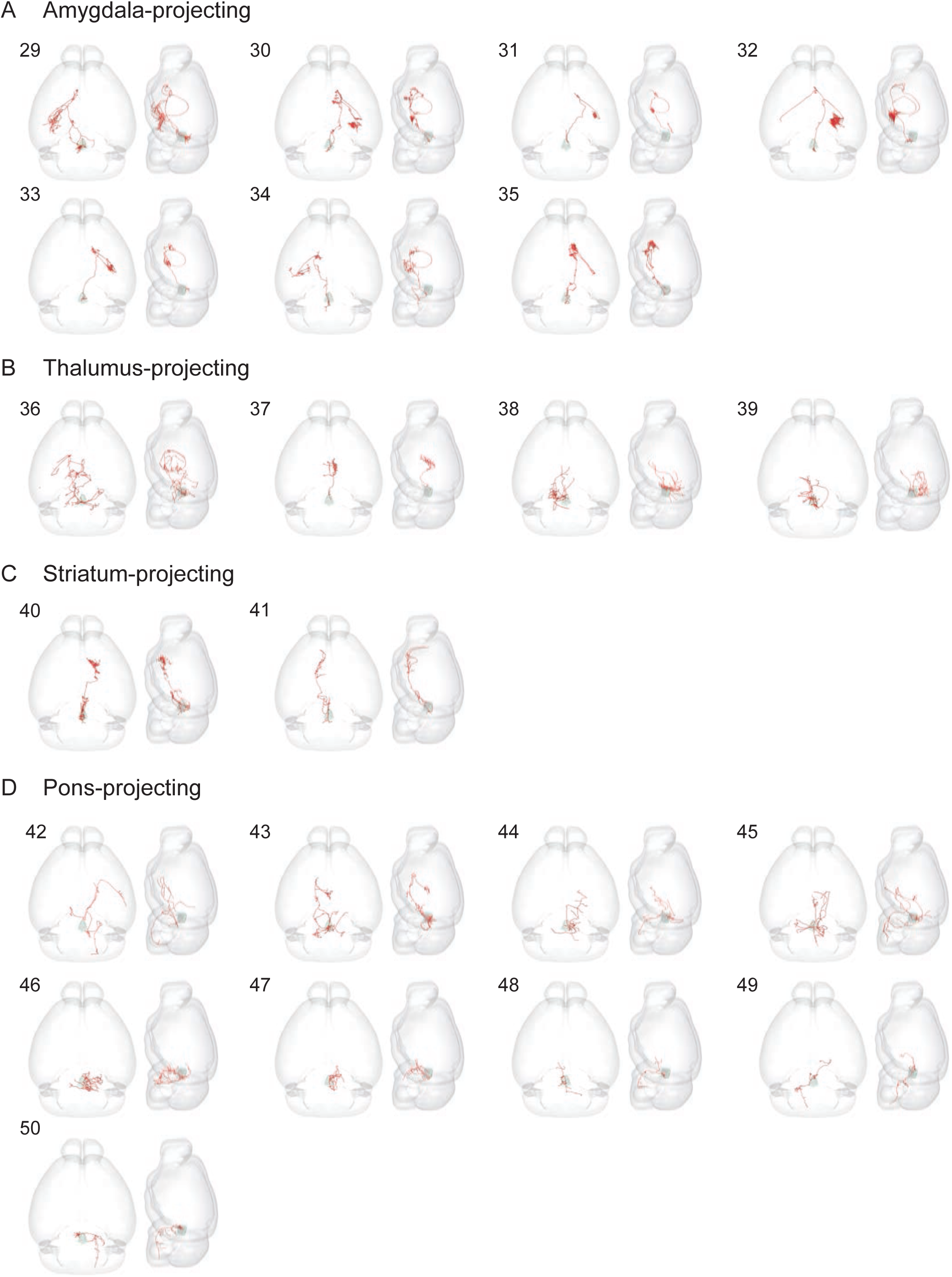
Individual reconstructed DR serotonin neurons (part II). Horizontal (left) and sagittal (right) view of 7 amygdala-projecting (**A**), 4 thalamus-projecting (**B**), 2 striatum-projecting (**C**), and 9 caudal brainstem-projecting (**D**) DR serotonin neurons.

**Figure 8—figure supplement 4.**
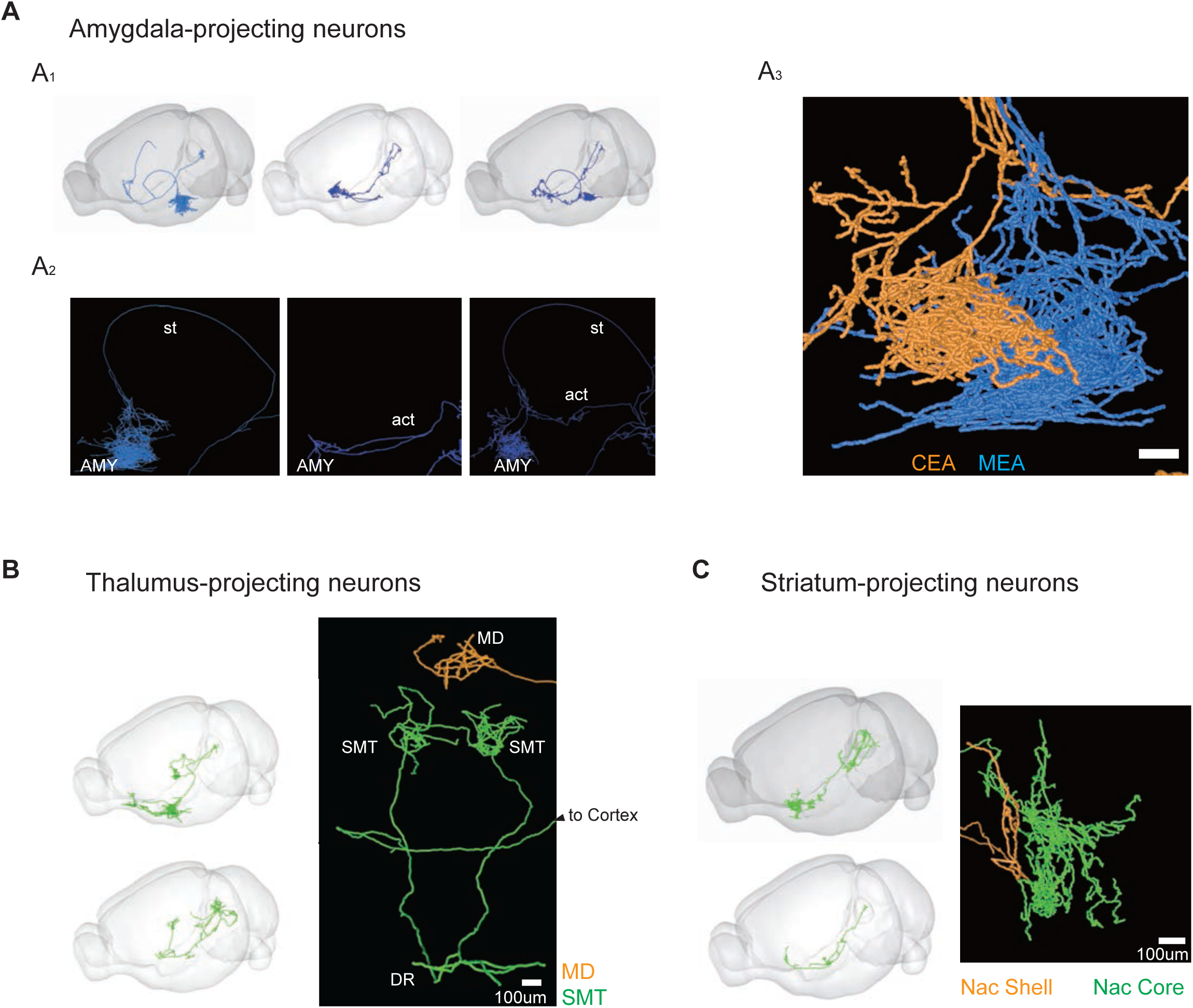
Detailed projection patterns of example DR serotonin neurons. **(A)** Projection patterns of the amygdala-projection serotonin neurons. Schematic (A1) and images (A2) of three types amygdala-projecting serotonin neurons. Some amygdala-projecting serotonin axons pass through the stria terminalis (left; n = 3), others pass through part of the anterior commissure (middle; n = 1), and yet others have branches that pass through both (right; n = 3). (A3) Examples of axon arbors of an amygdala-projecting serotonin neuron (yellow) that terminates in the central amygdala (CeA; yellow) and another one (blue) that terminates in the nearby medial amygdala (MeA). **(B)** Two examples of thalamus-projecting neurons. Top, one neuron projects to the medial dorsal nucleus of the thalamus (MD) without branches in the cortex. Bottom, one neuron has bilateral projections to the submedial nucleus of the thalamus (SMT) and also project to the cortex (not shown). **(C)** Two examples of striatum-projecting neurons differentially arborize at the core and shell of the nucleus accumbens (NAc).

**Video 1: Fly-through of aligned axonal projections and heatmaps of the *Trh^+^* and *Vglut3^+^* serotonin subpopulations.** Related to Figure 7.

Left, individual slice Z projection represent 125 um depth; Right, heat map of axonal densities calculated as described in Figure 7.

**Video 2: Whole-brain axonal projection patterns of 6 reconstructed dorsal raphe serotonin neurons.** Related to Figure 8.

Reconstructed serotonin neurons shown in Figure 8A–E were merged and presented in the standard brain, corresponding to Figure 8F.

**Supplemental Table 1. Raw data for single-cell gene expression.** Column names denote cell index, row names denote gene names.

**Supplemental Table 2. Functional gene categories, used to generate Figure 4 and Supplemental file 1.**

**Supplemental Table 3. Pearson correlation coefficient (*rp*) of pairwise correlation of gene expression across 999 cells.** Contains gene pairs with *rp>0.3, rp<–0.3*.

**Supplemental Table 4. Allen Brain Atlas IDs and Their Corresponding Names as Identified by the 2017 Common Coordinate Framework, Related to Figure 7.** Regions were selected prior to analysis such that areas defined by individual layers (e.g., cortical layers I–VI), cell identity, and anatomical cardinal directions are collapsed into their parent region. Individual normalized regional densities for each brain are aligned to the heat maps from Figure 7E.

**Supplemental File 1. Co-expression networks.**

Networks were constructed based on Pearson correlation coefficient (*r*p) of gene expression across all cells. Genes appear connected if *r*p >0.4. Edge width represents *r*p. Nodes are colored according to functional gene categories (***Materials and Methods***).

## References

Anders, S., Pyl, P.T., and Huber, W. (2015). HTSeq--a Python framework to work with high-throughput sequencing data. Bioinformatics 31, 166–169.

Bang, S.J., Jensen, P., Dymecki, S.M., and Commons, K.G. (2012). Projections and interconnections of genetically defined serotonin neurons in mice. Eur J Neurosci 35, 85–96.

Bedford, F.K., Julius, D., and Ingraham, H.A. (1998). Neuronal expression of the 5HT3 serotonin receptor gene requires nuclear factor 1 complexes. J Neurosci 18, 6186–6194.

Belmaker, R.H., and Agam, G. (2008). Major depressive disorder. N Engl J Med 358, 55–68.

Butler, A., Hoffman, P., Smibert, P., Papalexi, E., and Satija, R. (2018). Integrating single-cell transcriptomic data across different conditions, technologies, and species. Nat Biotechnol 36, 411–420.

Calizo, L.H., Akanwa, A., Ma, X., Pan, Y.Z., Lemos, J.C., Craige, C., Heemstra, L.A., and Beck, S.G. (2011). Raphe serotonin neurons are not homogenous: electrophysiological, morphological and neurochemical evidence. Neuropharmacology 61, 524–543.

Çiçek, Ö., Abdulkadir, A., Lienkamp, S.S., Brox, T., Ronneberger, O. (2016). 3D U-Net: Learning Dense Volumetric Segmentation from Sparse Annotation. In: Ourselin S., Joskowicz L., Sabuncu M., Unal G., Wells W. (eds) Medical Image Computing and Computer-Assisted Intervention – MICCAI 2016. MICCAI 2016. Lecture Notes in Computer Science, vol 9901. Springer, Cham.

Chi, J., Crane, A., Wu, Z., and Cohen, P. (2018). Adipo-Clear: A Tissue Clearing Method for Three-Dimensional Imaging of Adipose Tissue. J Vis Exp.

Choi, H.M.T., Schwarzkopf, M., Fornace, M.E., Acharya, A., Artavanis, G., Stegmaier, J., Cunha, A., and Pierce, N.A. (2018). Third-generation in situ hybridization chain reaction: multiplexed, quantitative, sensitive, versatile, robust. Development 145.

Cohen, J.Y., Amoroso, M.W., and Uchida, N. (2015). Serotonergic neurons signal reward and punishment on multiple timescales. Elife 4.

Commons, K.G. (2015). Two major network domains in the dorsal raphe nucleus. J Comp Neurol 523, 1488–1504.

Dahlstrom, A., and Fuxe, K. (1964). Localization of monoamines in the lower brain stem. Experientia 20, 398–399.

Darmanis, S., Sloan, S.A., Zhang, Y., Enge, M., Caneda, C., Shuer, L.M., Hayden Gephart, M.G., Barres, B.A., and Quake, S.R. (2015). A survey of human brain transcriptome diversity at the single cell level. Proc Natl Acad Sci U S A 112, 7285–7290.

Deneris, E., and Gaspar, P. (2018). Serotonin neuron development: shaping molecular and structural identities. Wiley Interdiscip Rev Dev Biol 7.

Dobin, A., Davis, C.A., Schlesinger, F., Drenkow, J., Zaleski, C., Jha, S., Batut, P., Chaisson, M., and Gingeras, T.R. (2013). STAR: ultrafast universal RNA-seq aligner. Bioinformatics 29, 15–21.

Erbel-Sieler, C., Dudley, C., Zhou, Y., Wu, X., Estill, S.J., Han, T., Diaz-Arrastia, R., Brunskill, E.W., Potter, S.S., and McKnight, S.L. (2004). Behavioral and regulatory abnormalities in mice deficient in the NPAS1 and NPAS3 transcription factors. Proc Natl Acad Sci U S A 101, 13648–13653.

Fenno, L.E., Mattis, J., Ramakrishnan, C., Hyun, M., Lee, S.Y., He, M., Tucciarone, J., Selimbeyoglu, A., Berndt, A., Grosenick, L., et al. (2014). Targeting cells with single vectors using multiple-feature Boolean logic. Nat Methods 11, 763–772.

Fernandez, S.P., Cauli, B., Cabezas, C., Muzerelle, A., Poncer, J.C., and Gaspar, P. (2016). Multiscale single-cell analysis reveals unique phenotypes of raphe 5-HT neurons projecting to the forebrain. Brain Struct Funct 221, 4007–4025.

Gilbert, T.L. (2018). The Allen Brain Atlas as a Resource for Teaching Undergraduate Neuroscience. J Undergrad Neurosci Educ 16, A261–A267.

Gong, H., Xu, D., Yuan, J., Li, X., Guo, C., Peng, J., Li, Y., Schwarz, L.A., Li, A., Hu, B., et al. (2016). High-throughput dual-colour precision imaging for brain-wide connectome with cytoarchitectonic landmarks at the cellular level. Nat Commun 7, 12142.

Gong, S., Doughty, M., Harbaugh, C.R., Cummins, A., Hatten, M.E., Heintz, N., and Gerfen, C.R. (2007). Targeting Cre recombinase to specific neuron populations with bacterial artificial chromosome constructs. J Neurosci 27, 9817–9823.

Hay-Schmidt, A. (2000). The evolution of the serotonergic nervous system. Proc Biol Sci 267, 1071–1079.

He, M., Tucciarone, J., Lee, S., Nigro, M.J., Kim, Y., Levine, J.M., Kelly, S.M., Krugikov, I., Wu, P., Chen, Y., et al. (2016). Strategies and Tools for Combinatorial Targeting of GABAergic Neurons in Mouse Cerebral Cortex. Neuron 91, 1228–1243.

Hendricks, T., Francis, N., Fyodorov, D., and Deneris, E.S. (1999). The ETS domain factor Pet-1 is an early and precise marker of central serotonin neurons and interacts with a conserved element in serotonergic genes. J Neurosci 19, 10348–10356.

Ishimura, K., Takeuchi, Y., Fujiwara, K., Tominaga, M., Yoshioka, H., and Sawada, T. (1988). Quantitative analysis of the distribution of serotonin-immunoreactive cell bodies in the mouse brain. Neurosci Lett 91, 265–270.

Jacobs, B.L., and Azmitia, E.C. (1992). Structure and function of the brain serotonin system. Physiol Rev 72, 165–229.

Jensen, P., Farago, A.F., Awatramani, R.B., Scott, M.M., Deneris, E.S., and Dymecki, S.M. (2008). Redefining the serotonergic system by genetic lineage. Nat Neurosci 11, 417–419.

Jin, Y., Dougherty, S.E., Wood, K., Sun, L., Cudmore, R.H., Abdalla, A., Kannan, G., Pletnikov, M., Hashemi, P., and Linden, D.J. (2016). Regrowth of Serotonin Axons in the Adult Mouse Brain Following Injury. Neuron 91, 748–762.

Kast, R.J., Wu, H.H., Williams, P., Gaspar, P., and Levitt, P. (2017). Specific Connectivity and Unique Molecular Identity of MET Receptor Tyrosine Kinase Expressing Serotonergic Neurons in the Caudal Dorsal Raphe Nuclei. ACS Chem Neurosci 8, 1053–1064.

Kim, J.C., Cook, M.N., Carey, M.R., Shen, C., Regehr, W.G., and Dymecki, S.M. (2009). Linking genetically defined neurons to behavior through a broadly applicable silencing allele. Neuron 63, 305–315.

Kiyasova, V., and Gaspar, P. (2011). Development of raphe serotonin neurons from specification to guidance. Eur J Neurosci 34, 1553–1562.

Li, H., Courtois, E.T., Sengupta, D., Tan, Y., Chen, K.H., Goh, J.J.L., Kong, S.L., Chua, C., Hon, L.K., Tan, W.S., et al. (2017a). Reference component analysis of single-cell transcriptomes elucidates cellular heterogeneity in human colorectal tumors. Nat Genet 49, 708–718.

Li, H., Horns, F., Wu, B., Xie, Q., Li, J., Li, T., Luginbuhl, D.J., Quake, S.R., and Luo, L. (2017b). Classifying Drosophila Olfactory Projection Neuron Subtypes by Single-Cell RNA Sequencing. Cell 171, 1206–1220 e1222.

Lin, R., Wang, R., Yuan, J., Feng, Q., Zhou, Y., Zeng, S., Ren, M., Jiang, S., Ni, H., Zhou, C., et al. (2018). Cell-type-specific and projection-specific brain-wide reconstruction of single neurons. Nat Methods 15, 1033–1036.

Liu, Z.X., Zhou, J.F., Li, Y., Hu, F., Lu, Y., Ma, M., Feng, Q.R., Zhang, J.E., Wang, D.Q., Zeng, J.W., et al. (2014). Dorsal Raphe Neurons Signal Reward through 5-HT and Glutamate. Neuron 81, 1360–1374.

Luo, L., Callaway, E.M., and Svoboda, K. (2018). Genetic Dissection of Neural Circuits: A Decade of Progress. Neuron 98, 865.

Maddaloni, G., Bertero, A., Pratelli, M., Barsotti, N., Boonstra, A., Giorgi, A., Migliarini, S., and Pasqualetti, M. (2017). Development of Serotonergic Fibers in the Post-Natal Mouse Brain. Front Cell Neurosci 11, 202.

Madisen, L., Zwingman, T.A., Sunkin, S.M., Oh, S.W., Zariwala, H.A., Gu, H., Ng, L.L., Palmiter, R.D., Hawrylycz, M.J., Jones, A.R., et al. (2010). A robust and high-throughput Cre reporting and characterization system for the whole mouse brain. Nat Neurosci 13, 133–140.

Marcinkiewcz, C.A., Mazzone, C.M., D’Agostino, G., Halladay, L.R., Hardaway, J.A., DiBerto, J.F., Navarro, M., Burnham, N., Cristiano, C., Dorrier, C.E., et al. (2016). Serotonin engages an anxiety and fear-promoting circuit in the extended amygdala. Nature 537, 97–101.

McDevitt, R.A., Tiran-Cappello, A., Shen, H., Balderas, I., Britt, J.P., Marino, R.A.M., Chung, S.L., Richie, C.T., Harvey, B.K., and Bonci, A. (2014). Serotonergic versus nonserotonergic dorsal raphe projection neurons: differential participation in reward circuitry. Cell Rep 8, 1857–1869.

Mickelsen, L.E., Bolisetty, M., Chimileski, B.R., Fujita, A., Beltrami, E.J., Costanzo, J.T., Naparstek, J.R., Robson, P., and Jackson, A.C. (2019). Single-cell transcriptomic analysis of the lateral hypothalamic area reveals molecularly distinct populations of inhibitory and excitatory neurons. Nat Neurosci 22, 642–656.

Mondal, K., Ramachandran, D., Patel, V.C., Hagen, K.R., Bose, P., Cutler, D.J., and Zwick, M.E. (2012). Excess variants in AFF2 detected by massively parallel sequencing of males with autism spectrum disorder. Hum Mol Genet 21, 4356–4364.

Niederkofler, V., Asher, T.E., Okaty, B.W., Rood, B.D., Narayan, A., Hwa, L.S., Beck, S.G., Miczek, K.A., and Dymecki, S.M. (2016). Identification of Serotonergic Neuronal Modules that Affect Aggressive Behavior. Cell Rep 17, 1934–1949.

Ogawa, S.K., Cohen, J.Y., Hwang, D., Uchida, N., and Watabe-Uchida, M. (2014). Organization of monosynaptic inputs to the serotonin and dopamine neuromodulatory systems. Cell Rep 8, 1105–1118.

Okaty, B.W., Commons, K.G., and Dymecki, S.M. (2019). Embracing diversity in the 5-HT neuronal system. Nat Rev Neurosci.

Okaty, B.W., Freret, M.E., Rood, B.D., Brust, R.D., Hennessy, M.L., deBairos, D., Kim, J.C., Cook, M.N., and Dymecki, S.M. (2015). Multi-Scale Molecular Deconstruction of the Serotonin Neuron System. Neuron 88, 774–791.

Picelli, S., Bjorklund, A.K., Faridani, O.R., Sagasser, S., Winberg, G., and Sandberg, R. (2013). Smart-seq2 for sensitive full-length transcriptome profiling in single cells. Nat Methods 10, 1096–1098.

Picelli, S., Faridani, O.R., Bjorklund, A.K., Winberg, G., Sagasser, S., and Sandberg, R. (2014). Full-length RNA-seq from single cells using Smart-seq2. Nat Protoc 9, 171–181.

Pollak Dorocic, I., Furth, D., Xuan, Y., Johansson, Y., Pozzi, L., Silberberg, G., Carlen, M., and Meletis, K. (2014). A whole-brain atlas of inputs to serotonergic neurons of the dorsal and median raphe nuclei. Neuron 83, 663–678.

Ravindran, L.N., and Stein, M.B. (2010). The pharmacologic treatment of anxiety disorders: a review of progress. J Clin Psychiatry 71, 839–854.

Ren, J., Friedmann, D., Xiong, J., Liu, C.D., Ferguson, B.R., Weerakkody, T., DeLoach, K.E., Ran, C., Pun, A., Sun, Y., et al. (2018). Anatomically Defined and Functionally Distinct Dorsal Raphe Serotonin Sub-systems. Cell 175, 472–487 e420.

Renier, N., Adams, E.L., Kirst, C., Wu, Z., Azevedo, R., Kohl, J., Autry, A.E., Kadiri, L., Umadevi Venkataraju, K., Zhou, Y., et al. (2016). Mapping of Brain Activity by Automated Volume Analysis of Immediate Early Genes. Cell 165, 1789–1802.

Rosenberg, A.B., Roco, C.M., Muscat, R.A., Kuchina, A., Sample, P., Yao, Z., Graybuck, L.T., Peeler, D.J., Mukherjee, S., Chen, W., et al. (2018). Single-cell profiling of the developing mouse brain and spinal cord with split-pool barcoding. Science 360, 176–182.

Saunders, A., Macosko, E.Z., Wysoker, A., Goldman, M., Krienen, F.M., de Rivera, H., Bien, E., Baum, M., Bortolin, L., Wang, S., et al. (2018). Molecular Diversity and Specializations among the Cells of the Adult Mouse Brain. Cell 174, 1015–1030 e1016.

Sengupta, A., Bocchio, M., Bannerman, D.M., Sharp, T., and Capogna, M. (2017). Control of Amygdala Circuits by 5-HT Neurons via 5-HT and Glutamate Cotransmission. J Neurosci 37, 1785–1796.

Shekhar, K., Lapan, S.W., Whitney, I.E., Tran, N.M., Macosko, E.Z., Kowalczyk, M., Adiconis, X., Levin, J.Z., Nemesh, J., Goldman, M., et al. (2016). Comprehensive Classification of Retinal Bipolar Neurons by Single-Cell Transcriptomics. Cell 166, 1308–1323 e1330.

Spaethling, J.M., Piel, D., Dueck, H., Buckley, P.T., Morris, J.F., Fisher, S.A., Lee, J., Sul, J.Y., Kim, J., Bartfai, T., et al. (2014). Serotonergic neuron regulation informed by in vivo single-cell transcriptomics. FASEB J 28, 771–780.

Steinbusch, H.W. (1981). Distribution of serotonin-immunoreactivity in the central nervous system of the rat-cell bodies and terminals. Neuroscience 6, 557–618.

Stensrud, M.J., Puchades, M., and Gundersen, V. (2014). GABA is localized in dopaminergic synaptic vesicles in the rodent striatum. Brain Struct Funct 219, 1901–1912.

Stuart, T., Butler, A., Hoffman, P., Hafemeister, C., Papalexi, E., Mauck, W.M., 3rd, Hao, Y., Stoeckius, M., Smibert, P., and Satija, R. (2019). Comprehensive Integration of Single-Cell Data. Cell 177, 1888–1902 e1821.

Tabula Muris, C., Overall, c., Logistical, c., Organ, c., processing, Library, p., sequencing, Computational data, a., Cell type, a., Writing, g., et al. (2018). Single-cell transcriptomics of 20 mouse organs creates a Tabula Muris. Nature 562, 367–372.

Tasic, B., Hippenmeyer, S., Wang, C., Gamboa, M., Zong, H., Chen-Tsai, Y., and Luo, L. (2011). Site-specific integrase-mediated transgenesis in mice via pronuclear injection. Proc Natl Acad Sci U S A 108, 7902–7907.

Tasic, B., Menon, V., Nguyen, T.N., Kim, T.K., Jarsky, T., Yao, Z., Levi, B., Gray, L.T., Sorensen, S.A., Dolbeare, T., et al. (2016). Adult mouse cortical cell taxonomy revealed by single cell transcriptomics. Nat Neurosci 19, 335–346.

Tasic, B., Yao, Z., Graybuck, L.T., Smith, K.A., Nguyen, T.N., Bertagnolli, D., Goldy, J., Garren, E., Economo, M.N., Viswanathan, S., et al. (2018). Shared and distinct transcriptomic cell types across neocortical areas. Nature 563, 72–78.

Teissier, A., Chemiakine, A., Inbar, B., Bagchi, S., Ray, R.S., Palmiter, R.D., Dymecki, S.M., Moore, H., and Ansorge, M.S. (2015). Activity of Raphe Serotonergic Neurons Controls Emotional Behaviors. Cell Rep 13, 1965–1976.

Wang, H.L., Zhang, S., Qi, J., Wang, H., Cachope, R., Mejias-Aponte, C.A., Gomez, J.A., Mateo-Semidey, G.E., Beaudoin, G.M.J., Paladini, C.A., et al. (2019). Dorsal Raphe Dual Serotonin-Glutamate Neurons Drive Reward by Establishing Excitatory Synapses on VTA Mesoaccumbens Dopamine Neurons. Cell Rep 26, 1128–1142 e1127.

Weissbourd, B., Ren, J., DeLoach, K.E., Guenthner, C.J., Miyamichi, K., and Luo, L. (2014). Presynaptic partners of dorsal raphe serotonergic and GABAergic neurons. Neuron 83, 645–662.

Welch, J.D., Kozareva, V., Ferreira, A., Vanderburg, C., Martin, C., and Macosko, E.Z. (2019). Single-Cell Multi-omic Integration Compares and Contrasts Features of Brain Cell Identity. Cell.

Wu, H., Williams, J., and Nathans, J. (2014). Complete morphologies of basal forebrain cholinergic neurons in the mouse. Elife 3, e02444.

Wylie, C.J., Hendricks, T.J., Zhang, B., Wang, L., Lu, P., Leahy, P., Fox, S., Maeno, H., and Deneris, E.S. (2010). Distinct transcriptomes define rostral and caudal serotonin neurons. J Neurosci 30, 670–684.

Zeisel, A., Hochgerner, H., Lonnerberg, P., Johnsson, A., Memic, F., van der Zwan, J., Haring, M., Braun, E., Borm, L.E., La Manno, G., et al. (2018). Molecular Architecture of the Mouse Nervous System. Cell 174, 999–1014 e1022.

Zeisel, A., Munoz-Manchado, A.B., Codeluppi, S., Lonnerberg, P., La Manno, G., Jureus, A., Marques, S., Munguba, H., He, L., Betsholtz, C., et al. (2015). Brain structure. Cell types in the mouse cortex and hippocampus revealed by single-cell RNA-seq. Science 347, 1138–1142.

